# Myeloperoxidase (MPO) exacerbates dengue-associated liver injury and contributes to disease pathogenesis in mouse models

**DOI:** 10.64898/2026.08.23.746568

**Authors:** Carla Bianca Luena Victorio, Andrew Teo, Shantanu Gupta, Arun Ganasarajah, Joanne Ong, Jasmine SK, Kíssila Rabelo, Luciana Lontro Alves, Carlos Alberto Basílio-de-Oliveira, Rodrigo Panno Basílio-de-Oliveira, Po Ying Chia, Heshan Kuruppu, Maneshka Karunananda, Damayanthi Idampitiya, Ananda Wijewickrama, Chandima Jeewandara, Neelika Malavige, Tsin Wen Yeo, Ann-Marie Chacko

## Abstract

Severe dengue can damage the liver through unestablished mechanisms. We investigated the role of myeloperoxidase (MPO), a neutrophil enzyme, in dengue through patients, fatal liver samples, and mouse infection models. Observations from two independent clinical cohorts revealed elevated plasma MPO levels in dengue and, in one cohort, MPO was further linked to liver injury markers during the critical phase of disease, whereas livers from dengue fatal cases revealed MPO build-up in the vicinity of CD177^+^ activated neutrophils. In mice, dengue led to MPO overexpression, oxidative damage, and broad activation of innate and systemic inflammatory pathways in livers. Blocking MPO activity alleviated these and improved survival in one model and delayed disease progression without preventing death in another. These findings establish MPO as a functional mediator of severe dengue-associated liver injury and inflammation, which warrants further preclinical investigation into its hepatic pathogenic mechanism and its validity as target for therapeutic intervention.

## Introduction

Dengue virus (DENV) infection is the most prevalent arboviral disease globally, with 53% of the global population at risk of dengue infection, a figure projected to rise to 63% by 2070^1,2^. Of the approximately 400 million annual infections, only a quarter are symptomatic^3^. Clinical manifestations range from self-limiting febrile illness to severe dengue, characterised by vascular leakage, shock and multi-organ dysfunction. Among affected organs, the liver is a major site of involvement. Elevations in the transaminases, alanine aminotransferase (ALT) and aspartate aminotransferase (AST), are observed in most patients, with the magnitude of elevation rising in parallel with disease severity^4–9^. Although acute liver failure is relatively uncommon, occurring in an estimated 0.3–1.1% of dengue patients, it carries a high mortality rate (20–60%)^8,10–12^; and early transaminase elevation during the febrile phase has been identified as a predictor of progression to severe disease^13–15^. Despite its clinical relevance, the mechanisms underlying dengue-associated liver injury remain incompletely defined.

DENV infection in the liver has also been well documented in the literature. Histopathological studies indicate that DENV infects both hepatocytes and Kupffer cells, accompanied by hepatic necrosis, steatosis and inflammatory cell infiltration^6,16–18^. Experimental models recapitulate key features of human disease, including elevated transaminases and immune cell infiltration in the liver^19,20^. While direct viral cytopathic effects contribute to tissue injury^21,22^, accumulating evidence suggests that host immune responses play a central role in mediating hepatic damage. Specifically, innate immune activation and leukocyte recruitment have been implicated in driving hepatic inflammatory pathology^23–25^.

While Kupffer cells dominate hepatic immunity at steady state, injury can trigger rapid neutrophil recruitment. Neutrophils are among the earliest responders to infection and are a major source of myeloperoxidase (MPO), a heme-containing enzyme that generates reactive oxygen species, including hypochlorous acid^26,27^. While these oxidants contribute to antimicrobial defence, they can also induce oxidative damage to cellular macromolecules and exacerbate tissue injury^23–25,28–30^. MPO has been implicated in inflammatory liver diseases, such as alcoholic hepatitis and non-alcoholic steatohepatitis (NASH)^26,31,32^, and identified as a potential biomarker of severe dengue in transcriptional and clinical studies^33–36^. Additionally oxidative stress was shown to induce liver damage in dengue^37^. Despite these associations, whether MPO functionally contributes to hepatic injury in dengue, or merely a consequence of liver injury, remains untested.

Here, we investigate the role of MPO in DENV infection through the lens of dengue-associated liver injury, integrating evidence across clinical cohorts, postmortem human liver tissue, and complementary experimental mouse models of severe dengue. The association of plasma MPO with disease severity and liver injury in dengue patients is examined using both longitudinal and cross-sectional sampling from Singapore and Sri Lanka. MPO expression in postmortem livers from fatal dengue cases in Brazil and mouse models of severe dengue is determined to provide spatial context of MPO expression juxtaposed to disease. Lastly, the functional contribution of MPO to disease and hepatic injury in dengue is evaluated using two structurally distinct pharmacological inhibitors in two independent mouse models of DENV infection. Together, these studies evaluate the contribution of MPO to dengue-associated liver injury and explore the therapeutic potential of MPO-directed intervention.

## Results

### Elevated MPO and liver injury markers in severe dengue patients from Singapore

We conducted prospective clinical studies to investigate the association between MPO and liver injury in human DENV infections. At baseline, SD patients exhibited higher prevalence of hypertension (90%) and longer hospitalization (7 days) compared with DwoS (36%; 4 days) and DWS (58 %; 5 days; **Table 1**). Among participants, serotypes 2 and 3 (DENV-2/3) were the most common detected in 15 individuals, while serotype 4 (DENV-4) was detected in one (*data not shown*). NS1 levels showed no association with disease severity (**Table 1**).

**Table 1.** Baseline characteristics of the Singapore dengue participants classified by 2009 Dengue guidelines.

| Variables | Controls<br>(n = 30) | DwoS<br>(n = 25) | DWS<br>(n = 24) | SD<br>(n = 10) | p<sup>a</sup> |
| --- | --- | --- | --- | --- | --- |
| Male (%) | 15 (50.0) | 16 (64.0) | 15 (62.5) | 6 (60) | 0.252 |
| Median age (IQR<sup>b</sup>) [range], years | 44 (32-59)<br>[23-75] | 40 (32-55)<br>[24-73] | 48 (33-61)<br>[22-80] | 61 (36-68) [24-83] | 0.07 |
| Median Body Mass Index (IQR), kg/m<sup>2</sup> | 24.3<br>(21.9-26.4) | 27.2<br>(22.9-30.7) | 25.9<br>(23.3-30.7) | 26.2<br>(23.5-28.7) | 0.392 |
| Median CCI (IQR) [range] | 0 (0-0) [0-2] | 0 (0-0) [0-3] | 1 (0-0) [0-3] | 1.5 (0-1) [0-5] | 0.072 |
| Diabetes Mellitus, n (%) | 3 (10) | 4 (16.0) | 8 (33.3) | 3 (30) | 0.423 |
| Hypertension, n (%) | 4 (13.3) | 9 (36.0) | 14 (58.3) | 9 (90) | **0.001** |
| Previous Dengue, n (%) | 3 (10) | 1 (4.0) | 2 (8.3) | 2 (20.0) | 0.132 |
| Median day of illness for febrile phase (IQR) | N.A.<sup>c</sup> | 4<br>(3-5) | 4<br>(3-5) | 5<br>(4-5) | 0.868 |
| Median day of illness for critical phase (IQR) | N.A. | 6<br>(5-7) | 6<br>(5-7) | 6<br>(5.5-7) | 0.277 |
| Median day of illness for late recovery phase (IQR) | N.A. | 20.5<br>(16-24.5) | 17.5<br>(15-21.5) | 18<br>(15-26) | 0.102 |
| Length of hospital stay (IQR), days | N.A. | 4 (3-5.5)<sup>d</sup> | 5 (4-6) | 7 (5-8) | **0.010** |
| **NS1 (ng/mL)** |  |  |  |  |  |
| Febrile phase | N.A. | 283<br>(239-488) | 219<br>(0-650) | 113<br>(0-226) | 0.4 |
| Critical phase | N.A. | 200<br>(0-357) | 23<br>(0-278) | 20<br>(0-151) | 0.6 |

<sup>a</sup> Kruskal-Wallis or chi-squared test/Fisher's exact for comparison of controls, dengue without warning signs (DwoS), dengue with warning signs (DWS), and severe dengue (SD) groups.
<sup>b</sup> IQR, interquartile range.
<sup>c</sup> N.A., not applicable.
<sup>d</sup> 24/25 participants with DwoS were admitted during dengue illness.

Regardless of disease phase (*i.e.,* “combined”), DwoS, DWS, and SD patients exhibited 1.6- to 1.7-fold higher plasma MPO levels compared to healthy controls (8.8 ng/mL), with highest recorded in SD patients (14.7 ng/mL) (**Fig. 1a; Table S1**). In SD patients, MPO levels peaked at febrile phase, earlier than in DwoS and DWS patients where MPO peaked at the critical phase (**Fig. 1a,b; Table S1; Table S2**). MPO levels at febrile phase in SD (21.6 ng/mL) were also 1.6-fold higher than DwoS (13.7 ng/mL; *p* = 0.004) and DWS (13.8 ng/mL; *p* = 0.01) patients (**Fig. 1a; Table S1**). Further, regardless of disease severity, plasma MPO at febrile and critical phase were higher than healthy controls and recovered dengue patients (**Fig. 1b; Table S2**).

**Figure 1.**
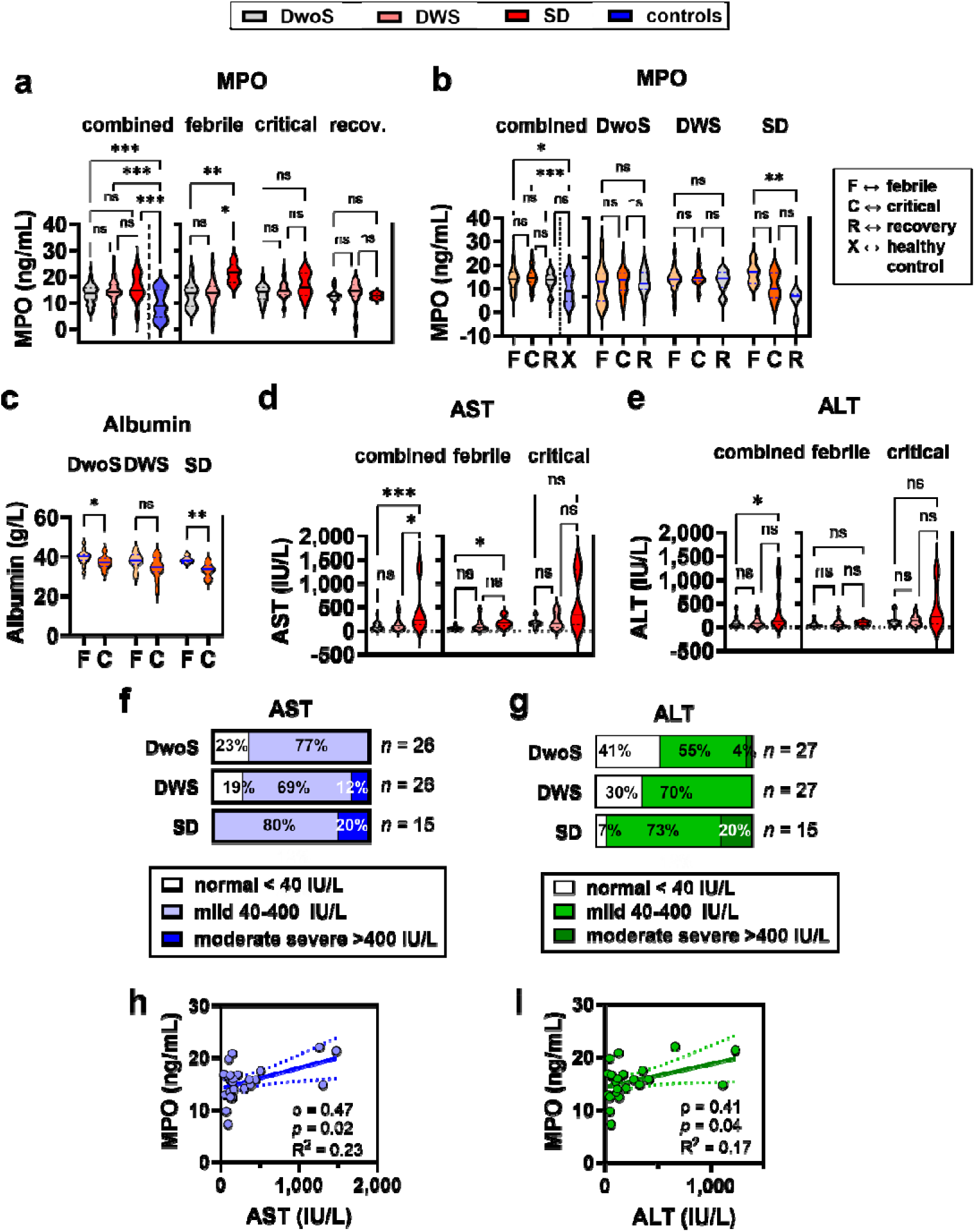
Circulating myeloperoxidase (MPO) is elevated in dengue and is associated with liver injury in the Singapore cohort. Dengue patients included in the trial were classified according to 2009 WHO criteria of disease severity as either dengue without warning signs (*DwoS*), dengue with warning signs (*DWS*), or severe dengue (*SD*). Healthy individuals (*controls*) were also recruited to determine baseline levels. Individuals were sampled multiple times during disease. **a-e,** Violin plots depicting median ± interquartile range (IQR) values of blood levels of MPO in dengue patients **a**, at febrile, critical, and recovery phases of diseases; or **b**, with varying disease severity; **c**, albumin; **d**, aspartate aminotransferase (AST); and **e**, alanine transaminase (ALT). Data in **a-e** are presented as median ± inter-quartile range (IQR) in bold lines and dashed lines, respectively, and medians in **a,b,d,e** were compared with Kruskal-Wallis test with Dunn’s correction for multiple comparisons. Medians in **c** were compared with Mann-Whitney test. f**-g**, Incidence of normal, mild, and moderate severe liver injury based on plasma levels of **f**, AST; and **g**, ALT. **h-i**, Monotonic correlation between plasma MPO and **h**, AST; or **i**, ALT. *ρ*, Spearman coefficient. *R^2^*, linearity coefficient. * *p* < 0.05. ** *p* < 0.005. *** *p* < 0.001. *ns*, not significant.

Despite neutropenia in disease relative to recovery (**Fig. S1a,b**), MPO-producing neutrophil counts at febrile and critical phase were highest in SD patients. At the critical phase of SD, absolute neutrophil counts (2,700 cells/μL) were 1.7-fold higher than in DwoS (1,560 cells/μL; *p* = 0.01) (**Table S1; Table S2**). Similar trends were observed for monocytes (**Fig. S1c,d**) but their levels were not modulated in SD patients (**Table S1; Table S2**).

Plasma albumin also declined from febrile to the critical phase, most evidently in SD, despite remaining within the normal range (34-54 g/L) in most patients. Albumin levels in SD at febrile phase (38 g/L, IQR 38-39) were 10.5% higher than at critical phase (34 g/L, IQR 30-35; *p* = 0.004) (**Fig. 1c**; **Table S1**) and 5% lower than in DwoS (40 g/L, IQR 38-41; *p* = 0.01) (**Fig. S2a; Table S1; Table S2**). These reductions reflect plasma leakage into tissues and/or reduced hepatic albumin synthesis consistent with hepatic involvement in SD patients.

More importantly, markers of liver injury—aspartate aminotransferase (AST) and alanine transaminase (ALT)—were also elevated from febrile to critical phase (**Fig. S2b,S2c**) and modulated by disease severity, with SD recording the highest levels (**Table S1; Table S2**). Regardless of disease phase, AST in SD patients (210 IU/L) was 3.6-fold higher than in DwoS (59 IU/L; *p* < 0.001) and 2.2-fold higher than in DWS (96 IU/L; *p* = 0.03), with comparable trends at febrile phase **(Fig. 1d**). Moreover, ALT in SD (110 IU/L) was 1.8-fold higher than DwoS (61 IU/L; *p* = 0.03) (**Fig. 1e**). Although no patient developed overt clinical hepatitis, the prevalence of mild (40–400 IU/L) and moderate-to-severe (> 400 IU/L) liver injury increased with disease severity (**Fig. 1f,g**).

Liver transaminases exhibited monotonic correlation with MPO. At the critical phase, MPO correlated with AST (Spearman, ρ = 0.47; *p* = 0.02) and ALT (ρ = 0.41; *p* = 0.04) regardless of disease severity (**Fig. 1h,i**). Such correlations were absent using data from the febrile phase alone, or at febrile and critical phases combined (**Fig. S3**).

### Elevated MPO and liver injury markers in severe dengue patients from Sri Lanka

We validated plasma MPO observations using retrospective data from an independent clinical cohort sampled at dengue febrile phase. At baseline, none of the primary comorbidities were associated with disease severity, although hypertension (11.4%) and diabetes mellitus (9.1%), were most prevalent in DwoS patients (**Table 2**). DENV-2 and DENV-3 were the most common strains detected in 42 and 10 patients, respectively, whilst serum NS1 was detected in 79% of patients (*data not shown*). Compared to the Singapore cohort, the median ages of study participants were lower in the Sri Lanka cohort—*e.g.* in SD, 61 years (IQR 36-68) *vs.* 30 years (IQR 21-34), respectively. Median body mass index (BMI) and prevalence of hypertension were also lower in the Sri Lanka cohort for DwoS (27.2 *vs*. 24.5; 36.0% *vs.* 11.4%), DWS (25.9 *vs*. 25.2; 58.3% *vs.* 3.7%), and SD (26.2 *vs*. 22.1; 90% *vs.* 0%) (**Table 1**; **Table 2**).

**Table 2.** Baseline characteristics of the Sri Lanka dengue participants classified by 2009 Dengue guidelines.

| **Variables** | **DwoS**<br>(n = 44) | **DWS**<br>(n = 27) | **SD**<br>(n = 7) | **p<sup>a</sup>** |
| --- | --- | --- | --- | --- |
| **Male (%)** | 28<br>(63.6) | 21<br>(77.8) | 5<br>(71.4) | 0.651 |
| **Median age (IQR) [range], years** | 36<br>(29-46) | 33<br>(26-45) | 30<br>(21-34) | 0.354 |
| **Median Body Mass Index (IQR), kg/m<sup>2</sup>** | 24.5<br>(20.7-26.9) | 25.2<br>(22.0-28.0) | 22.1<br>(19.5-23.2) | 0.36 |
| **Diabetes Mellitus, n (%)** | 4 (9.1) | 2 (7.4) | 0 (0) | 0.80 |
| **Hypertension, n (%)** | 5 (11.4) | 1 (3.7) | 0 (0) | 0.81 |

<sup>a</sup> Kruskal-Wallis or chi-squared test for comparison of dengue without warning signs (DwoS), dengue with warning signs (DWS), and severe dengue (SD) groups

Like the first cohort, MPO at febrile phase in the second cohort was modulated by disease severity (Kruskal-Wallis test, *p* < 0.001), but with the highest levels recorded in DWS (271.6 ng/mL, IQR 241.4-360.6) (**Table S3**). MPO in DWS was 1.4-fold higher than DwoS (197.7 ng/mL; *p* < 0.001) and 1.2-fold higher than SD (229.5 ng/mL; *p* = ns) (**Fig. 2a; Table S3**). Despite comparable leukocyte levels across groups, neutrophil counts were lowest in DWS (770 cells/µL), 35.8% lower than in DwoS (1,200 cells/µL; *p* < 0.001) and 46.2% lower than in SD (1,430 cells/µL; *p* = 0.02) (**Fig. 2b; Table S3**). Platelets were also reduced across all patients, with DWS and SD patients reaching severe thrombocytopenia (< 10^5^ platelets/µL). Compared to DwoS (10^5^ cells/µL), platelets declined by 70.3% in DWS (3×10^4^ cells/µL; *p* < 0.001) and by 55.4% in SD (4.5×10^4^ cells/µL; *p* = 0.002) (**Fig. 2c, Table S3**), suggesting higher haemorrhagic risk with increasing disease severity.

**Figure 2.**
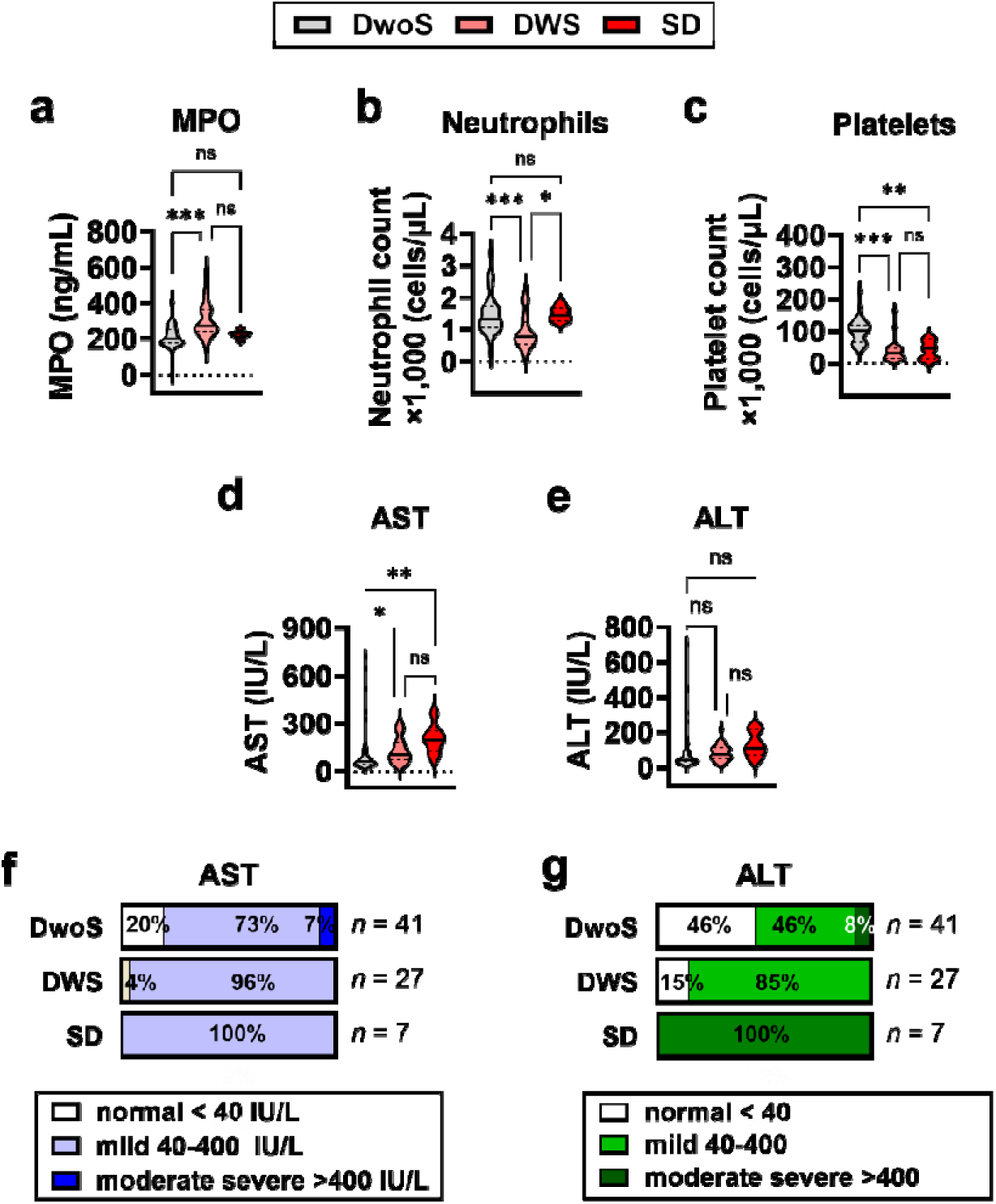
Circulating myeloperoxidase (MPO) is elevated alongside liver injury markers in the Sri Lanka dengue cohort. Dengue patients included in the trial were classified according to disease severity using the WHO 2009 criteria as either dengue without warning signs (*DwoS*), dengue with warning signs (*DWS*), or severe dengue (*SD*). **a-e,** Violin plots depicting median ± interquartile range (IQR) values in bold lines and dashed lines, respectively, for blood levels of **a**, MPO; **b**, neutrophils; **c**, platelet; **d**, aspartate aminotransferase (AST); and **e**, alanine transaminase (ALT). **f-g**, Incidence of normal, mild, and moderate severe liver injury based on plasma levels of **f**, AST; and **g**, ALT. Medians were compared with Kruskal-Wallis test with Dunn’s correction for multiple comparisons. * *p* < 0.05. ** *p* < 0.005. *** *p* < 0.001. *ns*, not significant.

Liver involvement was also evident with AST rising with increasing disease severity (*p* = 0.002), confirming observations from the first cohort (**Table S3**). Compared to DwoS (61 IU/L), AST was 1.7-fold higher in DWS (102 IU/L; *p* = 0.04) and 3.2-fold higher in SD (194 IU/L; *p* = 0.005) (**Fig. 2d, Table S3**). ALT exhibited a comparable trend (*p* = 0.03) but was not modulated by disease in pairwise comparisons (**Fig. 2e, Table S3**). Despite absence of overt hepatitis, mild and moderate severe liver injury were more prevalent in DWS and SD compared to DwoS (**Fig. 2f,g**). At febrile phase, plasma MPO and liver transaminase levels were not correlated, corroborating observations on febrile dengue patients in the Singapore cohort (**Fig. S4**).

The noted modulation of MPO and liver injury markers by disease severity was still evident even when patients in the second cohort was reclassified according to the 2011 WHO guidelines—dengue fever (DF), dengue haemorrhagic fever (DHF), and dengue shock syndrome (DSS). This was the default classification system used at the National Institute of Infectious Diseases, Sri Lanka (**Table S4**; **Fig. S5**). In DHF, plasma MPO (259.3 ng/mL) was 1.2-fold higher than in DF (220.5 ng/mL; *p* = 0.001); AST (139 IU/L *vs.* 63 IU/L; *p* = 0.004) and ALT (96 IU/L *vs.* 44 IU/L; *p* = 0.01) revealed similar trends (**Table S5**).

### *In situ* neutrophil activation and MPO accumulation in postmortem dengue liver

To confirm *in situ* MPO elevation in the liver, we conducted immunohistochemical (IHC) staining for MPO and CD177, a marker for activated neutrophils, on banked hepatic tissue sections from four confirmed dengue fatal cases from Brazil and a non-infectious control.

Control liver displayed minimal MPO staining, while dengue cases exhibited diffuse, intense MPO immunoreactivity concentrated within sinusoidal immune cells and extending into the hepatic parenchyma (**Fig 3a**, black arrows). Signal quantification (pixels per area of 0.96mm^2^) confirmed overt increased MPO levels within livers. Relative to control (103 pixels /area), MPO positivity was 25- to 29-fold higher in Case 1 (3,021 pixels/area; *p* < 0.001), Case 2 (2,536 pixels/area; *p* = 0.001), and Case 3 (2,707 pixels/area; *p* < 0.001) (**Fig. 3b; Table S6**). Case 4 exhibited only diffuse MPO signals (122 pixels/area, *p* = 0.8) (**Fig. 3b; Table S6**).

**Figure 3.**
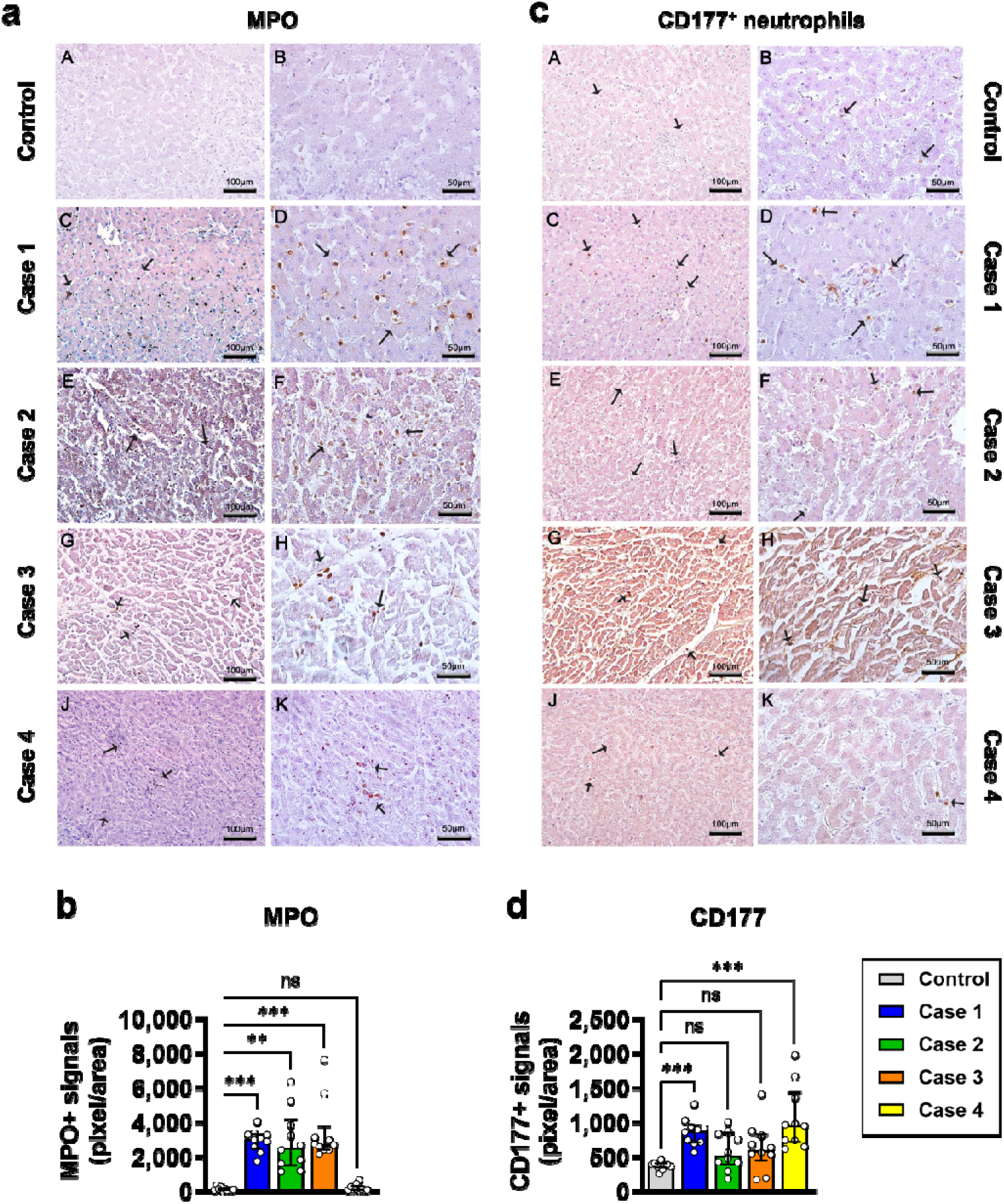
Myeloperoxidase (MPO)-positive activated neutrophils infiltrate livers from fatal dengue cases in Brazil. Human liver samples were collected from four fatal cases of dengue fever that occurred in Rio de Janeiro, Brazil. Control liver was taken from an individual who died of non-infectious causes and exhibited no evidence of hepatic pathology. **a-b,** MPO in liver samples. **a**, immunohistochemical (IHC) staining, where MPO-positive cells are highlighted (*black arrows*). **b**, Quantification of positive signals. **c-d**, Expression of CD177, a marker of activated neutrophils, in liver samples. **c**, immunohistochemical (IHC) staining, where CD177-positive cells are highlighted (*black arrows*). **d**, Quantification of positive signals in 10 fields of view (FOV) per section imaged. Each FOV is equivalent to 0.96 mm^2^, referred to as “*area*”. Data are shown as median ± interquartile range (IQR), and groups were compared by Kruskal-Wallis test with Dunn’s correction for multiple comparisons. * *p* < 0.05. ** *p* < 0.005. *** *p* < 0.001 *ns*, not significant.

Neutrophil identity was independently confirmed by CD177. Compared to control (376 pixels/area), CD177-positivity was markedly elevated 2.3- and 2.5-fold in Case 1 (875 pixels /area; *p* <0.001) and Case 4 (956 pixels /area; *p* <0.001), respectively, and were distributed throughout hepatic sinusoids and parenchyma, mirroring the MPO distribution (**Fig. 3c,d, Table S6**). Cases 2 and 3 exhibited insignificant increase in signals (1.38- and 1.59-fold *vs*. control) (**Fig. 3c,d, Table S6**).

### Elevated MPO and liver injury markers in mouse models of severe dengue

We next investigated whether clinical observations could be reproduced in two experimental DENV-2 infection systems using an antibody-dependent enhancement (ADE) model in A129 mice and a non-ADE severe dengue model in AG129 mice.

Immunohistochemical staining of livers at day 4 post-infection (p.i.) revealed increased CD45□ immune cell infiltration and MPO deposition, with MPO co-localising with Ly6G□ neutrophils, (**Fig. 4a,b**). Neutrophil activation was further confirmed by deposition of neutrophil elastase (**Fig. S6**) and corroborated postmortem findings in fatal clinical dengue.

**Figure 4.**
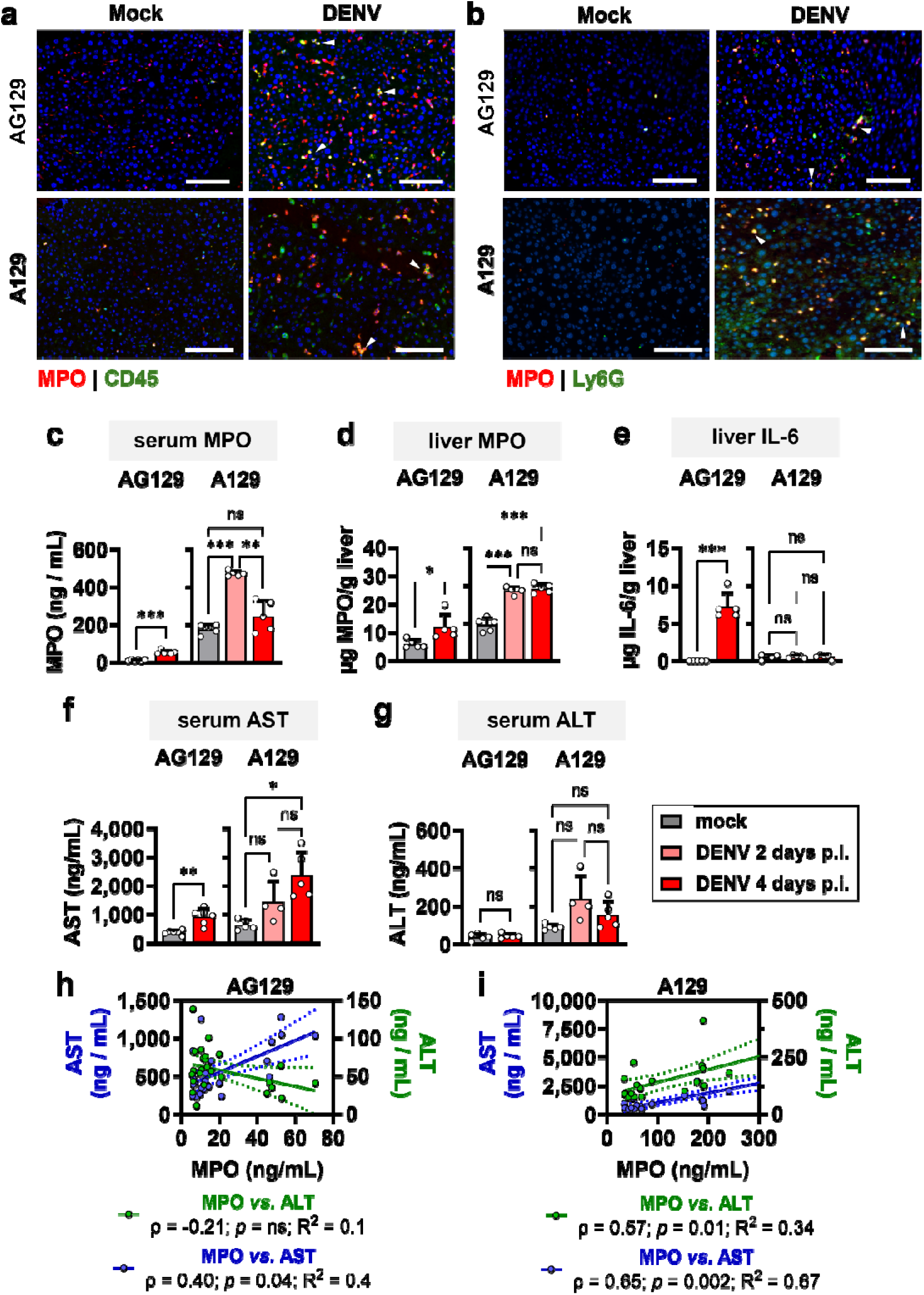
Myeloperoxidase (MPO) elevation marks neutrophil-rich liver inflammation and hepatic injury in dengue mouse models. AG129 severe dengue and A129 dengue ADE mouse models were evaluated at 2 and 4 days post infection (p.i.). **a-b,** Immunofluorescence staining of liver sections at 4 dpi depicting **a**, MPO in total CD45□ inflammatory cells; and **b**, MPO in Ly6G□ neutrophils. Colocalized signals are indicated by white arrows. **c–g,** Quantification of MPO levels in **c**, serum and **d**, liver homogenates; **e**, IL-6 levels in liver; and serum levels of **f**, aspartate aminotransferase (AST) and **g**, alanine transaminase (ALT). Data are presented as mean□±□SEM and representative of two independent studies. Statistical comparisons were performed in **c-g** using either two-tailed Welch’s *t*-test or Welch’s ANOVA with Dunnett’s correction for multiple comparisons. **h-i,** Monotonic correlation between serum MPO levels and AST or ALT levels in **h**, AG129 and **i**, A129 mice. _ρ_, Spearman’s correlation coefficient. *R^2^*, linearity coefficient. * *p*□<□0.05; ** *p*□<□0.005; *** *p*□<□0.001; *ns*, not significant.

Compared to mock-infected controls, serum MPO increased in AG129 mice at day 4 p.i. (5.4□±□1.0-fold higher; *p* < 0.001) but peaked at day 2 in A129 mice (2.7□±□0.1-fold higher; *p* < 0.001; **Fig. 4c**), with corresponding increases in MPO from liver homogenates (**Fig. 4d**). Notably, MPO in A129 mouse livers remained 1.2-fold higher than controls even at day 4 p.i. (*p* < 0.001; **Fig. 4d**). Both models also exhibited pronounced hepatic inflammation revealed by IL-6 overexpression in AG129 mice (>2,000-fold *vs.* controls; *p* < 0.001) (**Fig. 4e**) and broad upregulation of pro-inflammatory cytokines and chemokines in infected livers from both models (**Fig. S7**).

DENV infection also resulted in elevated markers of liver injury. Serum AST at day 4 p.i. was higher in AG129 (2.8□±□0.6-fold; *p* = 0.001) and A129 mice (3.7□±□1.3-fold; *p* = 0.03) than controls (**Fig. 4f**), although the increase was not reflected in liver homogenates (**Fig. S8**). In contrast, ALT was unchanged in both serum and liver tissue following infection (**Fig. 4g**; **Fig. S8**). Serum AST, but not ALT, exhibited monotonic correlation with serum MPO (_ρ_ = 0.40; *p* = 0.04) in AG129 mice (**Fig. 4h**). In A129 mice, both serum AST (_ρ_ = 0.65; *p* = 0.002) and ALT (_ρ_ = 0.57; *p* = 0.01) exhibited monotonic increase with serum MPO (**Fig. 4i**) and largely recapitulate observations from the longitudinal dengue cohort.

### MPO inhibition improves survival and reduces liver inflammation in the dengue ADE model

The functional contribution of MPO to dengue severity was further assessed by inhibiting MPO activity using two structurally distinct small-molecule inhibitors AZD5904 and mitiperstat (MTP) (**Fig. 5a**). Surprisingly in the A129 ADE model, both inhibitors markedly prolonged survival relative to untreated controls (median survival > 30 d *vs.* 5 d; *p* = 0.04 for AZD5904; *p* = 0.005 for MTP; **Fig. 5b**) and dampened infection-driven weight loss (**Fig. S9a**).

**Figure 5.**
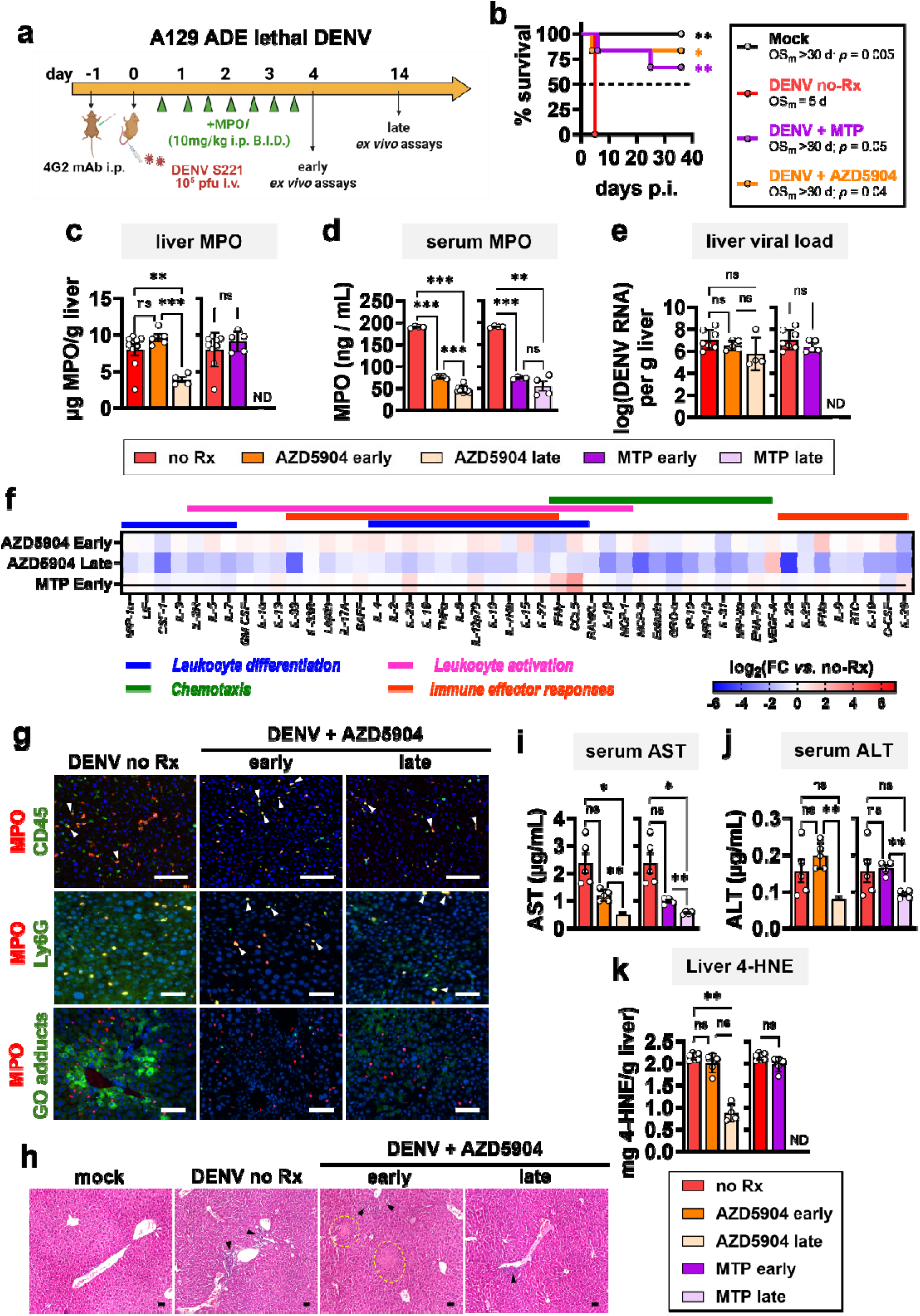
Myeloperoxidase (MPO) inhibition improves survival and attenuates DENV-associated liver injury in A129 mice. **a,** Study timeline. A129 mice were pre-injected with a high-affinity, non-neutralizing pan-flavivirus antibody and subsequently infected with dengue virus (DENV). Mice received twice-daily treatment with the MPO inhibitors (+MPO*i*) AZD5904 or mitiperstat (MTP); and were compared with infected mice receiving vehicle control (DENV no-Rx) and uninfected controls (Mock). Assays were conducted either at 4 days post-infection (p.i.; early) or 14 days p.i. (late). **b,** Kaplan–Meier survival curves following MPO*i* treatment. Survival was analysed using the Mantel–Cox test with Holm-Šídák’s correction for multiple comparisons *vs.* the DENV no-Rx group. **c–e,** Quantification of MPO levels in **c**, liver and **d**, serum; and **e**, viral load in liver. **f,** Heatmap of protein expression in liver homogenates relative to DENV no-Rx. Chemokines and cytokines involved in leukocyte migration (chemotaxis), differentiation, activation, and immune effector responses are shown. Data are presented as fold-change relative to DENV-infected, untreated mice (FC *vs.* no-Rx). **g**, Immunofluorescence staining of liver sections depicting expression of MPO, CD45 in total inflammatory cells, Ly6G in neutrophils, and guanine oxidation (GO) products. Colocalized signals are indicated by white arrows. **h**, Liver sections stained with hematoxylin and eosin. Inflammatory infiltrates are indicated by black arrows, and proteinaceous oedema are shown by green arrows. Regions of tissue necrosis are outlined in yellow dashed lines. **i-j**, Serum levels of **i**, aspartate aminotransferase (AST); and **j**, alanine transaminase (ALT). **k**, Lipid peroxidation assay measuring 4-hydroxynonenal (4-HNE) in liver homogenates. Data in **c-e** and **i-k** are presented as mean□±□SEM, representative of two independent studies. Means between two groups were compared by Welch’s *t*-test; means from > 2 groups were compared using either ordinary one-way ANOVA with Tukey’s correction, or Welch’s ANOVA with Dunnett’s correction, for multiple comparisons. * *p*□<□0.05; ** *p*□<□0.005; *** *p*□<□0.001; *ns*, not significant. *ND*, not determined. Scale bars = 50 µm.

Target engagement was also confirmed biochemically: AZD5904 reduced hepatic MPO at late disease by 50% (±□10%; *p* = 0.003) and plasma MPO by 75% (±□4%; *p* < 0.001) relative to untreated mice, with comparable effects for MTP (**Fig. 5c,d**). Neither inhibitor abrogated viral replication, producing only a modest reduction in liver viral load (0.8–0.9 log; **Fig. 5e**), indicating that the survival benefit was largely uncoupled from direct antiviral activity.

MPO inhibition broadly attenuated hepatic inflammation, most prominently at later timepoints. Affected mediators spanned myeloid maturation (CSF-1, IL-19), chemotaxis (MCP-1, MCP-3, GRO-α, ENA-78), and adaptive immune activation (IL-2R, IL-2, IL-4) (**Fig. 5f**). By day 14 p.i. (late treatment), AZD5904 suppressed regulators of neutrophil activity (MIP-1β, MIP-1α, LIF, eotaxin) and master systemic inflammatory cytokines (IL-6, IL-17, TNF-α) (**Fig. S10a**); MTP treatment resulted in more modest reductions in these same mediators (**Fig. S10b**). Neither inhibitor substantially altered the inflammatory milieu at early timepoint (day 4 p.i.; **Fig. 5f**; **Fig. S10**).

Consistent with reduced inflammation, treated livers exhibited diminished immune cell infiltration, particularly of Ly6G□ neutrophils and markedly lower *in situ* MPO signal, corroborating ELISA findings (**Fig. 5c,d,g**).

### MPO inhibition reduced oxidative damage to hepatic nucleic acids and reduced livery injury

Canonical guanine-oxidation (GO) products were detected in infected livers compared to healthy tissue, and MPO inhibition reduced their formation (**Fig. 5g**; **Fig. S11a**). Consistently, histological analysis revealed reduced inflammatory infiltration, interstitial oedema, and tissue necrosis after MPO inhibition (**Fig. 5h**; **Fig. S11b**).

These improvements in tissue integrity were accompanied by reduced biochemical liver injury. At late treatment timepoint with AZD5904, serum AST decreased by 79% (±□3.5%; *p* = 0.02) and 58% (±□6.9%; *p* = 0.003) *vs.* no treatment and early treatment timepoints, respectively; with similar outcomes following MTP treatment (**Fig. 5i**). Similarly, treatment with AZD5904 resulted in 61% (±□2.6%; *p* = 0.005) reduction in serum ALT at late *vs.* early timepoint, while MTP resulted in 44% (±□7.2%; *p* = 0.004) reduction (**Fig. 5j**). However, these changes were less apparent in liver homogenates (**Fig. S11c**).

More importantly, MPO inhibition diminished lipid peroxidation in livers, most evidently at late timepoint post-treatment with AZD5904 where 4-hydroxynonenal (4-HNE) levels (868 µg/g tissue) were down by 59.6% (*p* < 0.001) and 56.5% (*p* < 0.001) compared to healthy control (2,147 µg/g tissue) and treated liver at early timepoint (1,996 µg/g tissue), respectively (**Fig. 5k**). MTP treatment similarly lowered 4-HNE by 8.3% (*p* = 0.03) compared to healthy livers at early timepoint (**Fig. 5k**). Interestingly, liver MPO directly correlated with 4-HNE levels (ρ = 0.44, *p* = 0.014) but not with liver transaminases (**Fig. S12**).

### MPO inhibition delays disease progression and dampens liver inflammation in the AG129 model of severe dengue

The effect of MPO inhibition in DENV infection was also investigated in the more stringent AG129 model of lethal dengue (**Fig. 6a**). Low dose inhibitor (10 mg/kg) conferred no survival benefit (*data not shown*), whereas the higher dose (50 mg/kg) modestly extended survival (median 6-7 d *vs.* 4 d; *p* = 0.02 with MTP; *p* = 0.03 with AZD5904) and alleviated weight loss without preventing mortality (**Fig. 6b; Fig. S9b**). Both inhibitors reduced MPO levels in the circulation but not the liver (**Fig. 6c,d**). AZD5904 lowered mean plasma MPO by 27% (± 4.2%; *p* = 0.03) and 56% (± 12.0%; *p* < 0.001) at early and late timepoints, respectively; and MTP reduced it by 43% (± 14.6%; *p* < 0.001) and 60% (± 20.1%; *p* < 0.001) (**Fig. 6c**). As in the ADE model, MPO inhibition produced only a modest reduction in viral loads (0.7-0.9 log; *p* < 0.001; **Fig. 6e**), again indicating that any benefit was not primarily a result of direct antiviral activity.

**Figure 6.**
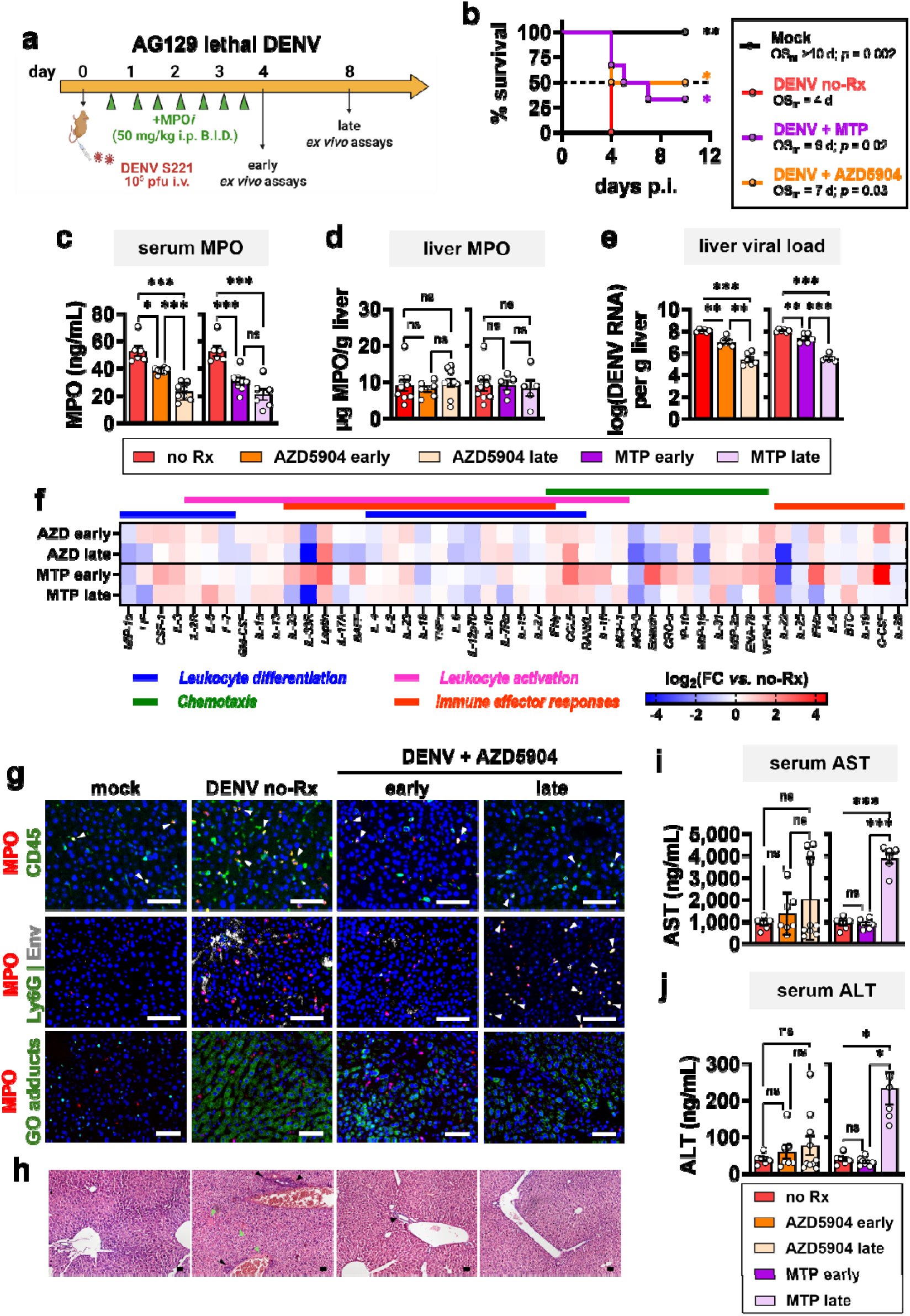
Myeloperoxidase (MPO) inhibition delays disease progression and attenuates hepatic inflammation and oxidative injury in AG129 mice. **a,** Study timeline. AG129 mice were infected with dengue virus (DENV), received twice-daily treatment with the MPO inhibitors (+MPO*i*) AZD5904 or mitiperstat (MTP); and were compared with infected mice receiving vehicle control (DENV no-Rx) and uninfected controls (Mock). Assays were conducted either at 4 days post-infection (p.i.; early) or 8 days p.i. (late). **b,** Kaplan–Meier survival curves following MPO*i* treatment. Survival was analysed using the Mantel–Cox test with Holm-Šídák’s correction for multiple comparisons *vs.* the DENV no-Rx group. **c–e,** Quantification of MPO levels in **c**, serum and **d**, liver homogenates; and **e**, viral load in liver. **f,** Heatmap of protein expression in liver homogenates relative to DENV no-Rx. Chemokines and cytokines involved in leukocyte migration (chemotaxis), differentiation, activation, and immune effector responses are shown. Data are presented as fold-change relative to DENV-infected, untreated mice (FC *vs.* no-Rx). **g**, Immunofluorescence staining of liver sections depicting expression of MPO, CD45 in total inflammatory cells, Ly6G in neutrophils, and guanine oxidation (GO) products. Colocalized signals are indicated by white arrows. **h**, Liver sections stained with hematoxylin and eosin. Inflammatory infiltrates are indicated by black arrows, and proteinaceous oedema are shown by green arrows. Regions of tissue necrosis are outlined in yellow dashed lines. **i-j**, Serum levels of **i**, aspartate aminotransferase (AST); and **j**, alanine transaminase (ALT). Data in **c-e** and **i-j** are presented as mean□±□SEM, representative of two independent studies. Means were compared using either ordinary one-way ANOVA with Tukey’s correction, or Welch’s ANOVA with Dunnett’s correction, for multiple comparisons. * *p*□<□0.05; ** *p*□<□0.005; *** *p*□<□0.001; *ns*, not significant. Scale bars = 50 µm.

MPO inhibition also broadly attenuated hepatic inflammatory responses (**Fig. 6f**). AZD5904 strongly suppressed cytokines involved in neutrophil maturation and migration (GRO-α, LIF, MIP-1α, and MIP-1β), whereas MTP acted most potently on LIF and IFN-α (**Fig. 6f; Fig. S13**). For systemic mediators, AZD5904 reduced IL-17A with more modest effects on IL-6, IL-17E and IL-18, while MTP most strongly suppressed IL-18, particularly at early timepoints (**Fig. 6f; Fig. S13**).

Histological and immunofluorescence analyses revealed reduced infiltration of immune cells, particularly neutrophils, and less detected GO products at early timepoint in livers following MPO inhibition, though inflammatory markers and tissue injury re-emerged at later stages (**Fig. 6g,h**; **Fig. S14a,b**). Liver transaminases in serum and liver homogenates were not significantly reduced (**Fig. 6i,j; Fig. S14c,d**).

## Discussion

This study investigated the role of myeloperoxidase (MPO) in dengue-associated liver injury. Across two independent clinical dengue cohorts, postmortem human tissues, and two complementary dengue mouse models, MPO was found associated with dengue severity and liver injury. More importantly, MPO inhibition in two experimental DENV infection models demonstrated the functional contribution of MPO to inflammatory pathology, particularly in the liver. Observations from the longitudinal dengue cohort revealed MPO and liver transaminases increased with dengue severity throughout the disease. At the critical phase, MPO even correlated with liver injury markers (AST and ALT). The early peak of MPO at febrile phase of SD, in contrast to its later peak at critical phase in milder dengue, warrants further investigation as a potential early predictor of severe disease. Observations from the cross-sectional validation cohort at febrile phase revealed increased MPO with disease, but peaking in DWS instead of SD. Nevertheless, liver transaminases in the validation cohort were also elevated in SD compared to milder disease and consistent with previous reports^9,13,37^. The lack of correlation between MPO and liver transaminases at febrile phase corroborated results from the longitudinal cohort and highlight the value of critical phase as the point of convergence between MPO elevation and hepatic injury, which warrants more extensive longitudinal investigation in larger clinical cohorts.

At the tissue level, livers from fatal dengue cases accumulated MPO and activated neutrophils. This feature was corroborated in mouse models, with accompanying increases in MPO, hepatic inflammation, and liver transaminase elevation. The most compelling evidence supporting a functional role for MPO contributing to dengue-associated pathology came from intervention experiments. Across two severe dengue models, pharmacological inhibition of MPO with either AZD5904 or mitiperstat (MTP) consistently attenuated inflammatory and oxidative injury in the liver and improved disease outcomes. Reduction of GO products—comprising of 8-hydroxy-2′-deoxyguanosine (oxo8dG, reporting DNA oxidation), 8-oxo-7,8-dihydroguanosine (oxo8G, reporting RNA oxidation), and the free base 8-oxo-7,8-dihydroguanine (oxo8Gua, reflecting oxidised nucleotides and repair turnover)—and decrease in 4-HNE in livers of mice treated with MPO inhibitors demonstrated lower oxidative burden on cellular nucleic acids and lipids. These were consistent with reduced differentiation and activation of neutrophils, as well as dampened vascular leakage, in DENV-infected AG129 mice treated with MPO inhibitor ABAH^38^. The consistent MPO elevation caused by DENV infection and reproducible effect of MPO inhibition across independent dengue models further solidify MPO as a biologically relevant contributor to dengue-associated pathology rather than simply a model-specific finding. These also provide additional impetus for further investigations into MPO-directed therapeutic interventions in dengue. Notably, the reductions in viral load achieved with MPO inhibition were well below the potency of well-studied DENV replication inhibitors^39^, indicating that the beneficial effect of MPO inhibition was achieved through a host-directed, rather than virus-targeted, mechanism.

Mechanistically, our findings were consistent with a model in which neutrophil-derived MPO contributes to hepatic injury through oxidative and inflammatory pathways. MPO potently mediates tissue oxidative stress by catalyzing the generation of hypochlorous acid (HOCl) and production of reactive oxygen species (ROS) and reactive nitrogen species (RNS) during the respiratory burst of activated neutrophils and monocytes. Conversely, MPO inhibition reduces HOCl production and oxidative stress^40^, thereby lessening tissue injury and improving outcomes observed in our experimental dengue models. DENV infection leads to increased hepatic infiltration of neutrophils, which produce RNS and ROS that oxidize cellular macromolecules, damage hepatocytes^23–30^, and promote cell death pathways such as apoptosis and necroptosis^41^. GO adducts of damaged nucleic acids in DENV infection provide a stable molecular footprint of this activity. Its reduction following MPO inhibition provides a mechanistic link between MPO activity and oxidative hepatocellular injury distinct from, and complementary to, the anti-inflammatory effects described below.

Secondly, MPO inhibition results in gross suppression of multiple inflammatory pathways suggesting an upstream regulatory role for MPO, although the precise position of MPO within these networks cannot be surmised from the present study. Rather than affecting isolated mediators, MPO inhibition suppressed three key inflammatory circuits in the liver: neutrophil recruitment, systemic cytokine amplification, and tissue-damaging immune signalling. Treatment with MPO inhibitors reduced hepatic expression of neutrophil chemotactic factors (*e.g.* MCP-1), master inflammatory switches (*e.g.* IL-6), and regulators of tissue injury (*e.g*. IL-17/IL-33 axes). Prior studies suggested MPO influences the recruitment, differentiation, activation, and downstream inflammatory signalling of neutrophils ^38,42,43^. MPO was shown to amplify a neutrophil–IL-17 inflammatory loop linked to elevated liver transaminase levels in dengue patients and shown to exacerbate hepatic damage in murine models^9,44,45^. The more pronounced cytokine changes at late treatment—*i.e.,* a week after the last drug dose, *vs.* early treatment further demonstrates the lasting effect of MPO inhibition on dampening the inflammatory responses.

Lastly, other proteins could have played a role in exacerbating liver injury following infection. DENV non-structural proteins are known to amplify MPO-mediated injury by depleting glutathione reserves and by promoting neutrophil extracellular trap (NET) formation^46^; thus creating a self-perpetuating cycle of inflammation correlated with disease severity^24,25,30^. Moreover, the proteolytic activity of MPO is known to exacerbate injury to hepatic connective tissue *via* modulation of matrix metalloproteinases (MMP),^28^ highlighting its multifaceted role in dengue-associated hepatitis.

Several study caveats are worth noting. First, epidemiologic variations between the two clinical cohorts at baseline, such as body mass index (BMI), prevalence of hypertension, timing of sampling, and low number of cases classified as severe dengue (SD), may have contributed to different trends in plasma MPO levels at febrile phase, thus restricting direct cross-cohort comparability. The clinical findings should be interpreted as evidence of association rather than proof of a causal relationship between MPO and liver injury. The observational nature of the human studies precludes causal inference. Second, postmortem liver analyses were restricted to a small number of fatal dengue cases where the proximity of MPO and CD177 suggested intrahepatic MPO were released by activated neutrophils rather than passive protein deposition. Though it adequately provided spatial evidence of MPO and neutrophil accumulation in the liver, it couldn’t establish temporal causality. Third, the mouse models used here recapitulate key features of severe dengue and dengue-associated liver inflammation, but their interferon-deficient background, viral challenge conditions, and antibody-enhancement setting could not fully capture the complexity of human dengue pathogenesis. Finally, while MPO inhibition substantiated the functional involvement of MPO in experimental disease, off-target drug effects could not be completely excluded. These experiments did not investigate contributions from additional neutrophil-derived mediators or broader effects of MPO inhibition on immune cell recruitment and activation. Further studies in larger, temporally harmonized patient cohorts and complementary immunocompetent preclinical models would be required to define the clinical window, safety, and translational potential of MPO-targeted therapy in dengue.

Collectively, our findings strongly support a role for MPO in dengue-associated pathology especially in the liver. Current management of acute liver failure in dengue lacks approved therapeutic strategies, with limited evidence supporting the increasing use of *N*-acetylcysteine as an adjuvant to manage severe dengue-induced hepatitis^47^. Given the beneficial effects of MPO inhibition in our experimental preclinical models, MPO inhibitors, particularly those that have entered human clinical development in other disease settings^48–50^, merit further investigation as potential therapeutic candidates. However, additional preclinical and clinical studies will be required to define efficacy, safety, optimal timing, and translational relevance in dengue.

## Online Methods

### Ethics statement

The clinical study in Singapore was approved by the National Healthcare Group Domain Specific Review Board (E/2016/00982). The clinical study in Sri Lanka was approved by the Ethics Review Committee of the Faculty of Medical Sciences, University of Sri Jayewardenepura, Sri Lanka (Ethics Application Number: 58/19). Informed written consent was obtained from all participants before enrolment in the study, and details of both study cohorts have been previously described^36,37^.

All experimental procedures and protocols involving fatal dengue cases and control samples were reviewed and approved by the Ethics Committee of the Pedro Ernesto University Hospital- Universidade do Estado do Rio de Janeiro (UERJ), Brazil under protocol number CAEE: 93175625.4.0000.5259. Written informed consent was obtained from families of participants.

### Clinical study details

Dengue subjects in Singapore were enrolled in prospective longitudinal study between September 2017 and December 2019. Participants were recruited during the febrile or critical phase, and samples were collected multiple times throughout hospitalization. Further, patients were followed up and samples collected again at recovery, 21–28 days post-discharge. DENV infection was confirmed *via* a positive NS1 antigen test (SD Bioline Dengue Duo, Korea), and pregnant or breastfeeding individuals were excluded. Controls consisted of adults with no febrile episodes in the two weeks prior to recruitment and no history of dengue infection in the preceding six months. See **Supplemental Methods** for full study details.

Of the 577 adult patients recruited to the previously reported DENV study at the National Institute of Infectious Diseases, Sri Lanka^37^, samples and blinded clinical data from seventy-eight (78) patients were included in this study. Patients were recruited between December 2022 and December 2024 during the febrile phase (within 4 days of illness onset). All those with known chronic kidney disease or chronic liver disease were excluded from the study. Plasma levels of transaminases and MPO were measured on the day of hospital admission.

DENV severity was classified using 2009 World Health Organization (WHO) criteria: dengue without warning signs (DwoS), dengue fever with warning signs (DWS), and severe dengue (SD)^51^. Typical warning signs include abdominal pain or tenderness, persistent vomiting, clinical fluid accumulation (ascites, pleural effusion), mucosal bleeding, liver enlargement, and unusual laboratory findings such as rising hematocrit with rapid platelet drop. Severe dengue was characterized by severe bleeding or severe plasma leakage leading to shock, as well as significant organ involvement.

### Blood Measurements and serological assays

Neutrophil, monocyte, and/or platelet counts, as well as liver transaminase and albumin levels, were analyzed by Tan Tock Seng Hospital Clinical Pathology Laboratory, Singapore. In the Sri Lankan cohort, these parameters were measured at the National Institute of Infectious Diseases. Blood samples were collected in lithium heparin or sodium citrate tubes, processed within 45 min of collection *via* centrifugation, and stored at -80 °C for plasma analysis. Plasma myeloperoxidase (MPO) was quantified using ELISA (Human Myeloperoxidase DuoSet ELISA, R&D Systems). Recombinant DENV-2 NS1 (baculovirus-derived; provided by Drs. Kitti Chan and Satoru Watanabe, Duke-NUS Medical School, Singapore) served as the standard for NS1 quantification *via* ELISA (Bio-Rad Platelia Dengue NS1 kit) as described previously^52^.

DENV serotyping and cycle threshold analysis were performed using the FDA-approved CDC DENV1-4 RT-PCR assay. Briefly, viral RNA was extracted from initial plasma samples and amplified in 25 µL singleplex reactions, following manufacturer instructions, containing 2 µL of RNA template.

### Immunohistochemical staining and imaging of liver samples from fatal dengue cases

Banked formalin-fixed paraffin-embedded (FFPE) tissue blocks of human liver tissue samples were previously obtained from four fatal cases of dengue fever that occurred in Rio de Janeiro, Brazil. All patients died with a clinical diagnosis of dengue hemorrhagic fever with classical symptoms (fever, myalgia and hemorrhagic manifestations) with DENV IgM-positive sera at hospital admission. Three of these cases (Cases 2–4) were associated with the severe dengue outbreak in 2002, predominantly caused by DENV-3^53^, while one case (Case 1) corresponded to the 2008 epidemic attributed to DENV-2^54^. FFPE liver sections used in this study were positive for DENV antigens and exhibited significant hemorrhaging, edema, and steatosis in immunohistopathological examinations^55,56^. Full details of the clinical history of the samples, as well as pathological findings on liver histology, were reported previously^55,56^ and briefly described in the **Supplemental Methods**. As a negative control, liver tissue was obtained from an individual who died of noninfectious causes and exhibited no evidence of hepatic pathology.

Formalin-fixed, paraffin-embedded (FFPE) liver sections were probed with the following primary antibodies: rat anti-human MPO monoclonal antibody (mAb) (Abcam, AB300650; 1:3000) and rabbit anti-human CD177 mAb (Abcam, AB255296; 1:500); and their corresponding horseradish peroxidase (HRP)-labelled secondary antibodies. HRP was detected with diaminobenzidine (DAB; Vector Laboratories, USA). Peroxidase staining was examined under a light microscope equipped with a CCD camera (Olympus BX53 with DP72 camera, Japan). Images from ten randomly selected, non-overlapping fields were acquired and analyzed using the Image-Pro Plus 7.0 software (Media Cybernetics, USA). Each field was 0.96 mm^2^ in area. All image acquisitions and analyses were performed by a single investigator blinded to the clinical diagnosis of the tissue samples to minimize observer bias.

### Mouse DENV infections

All animal procedures were conducted in accordance with National Institutes of Health (NIH) guidelines with approval from the SingHealth/Duke-NUS Institutional Animal Care and Use Committee (IACUC) (2020/SHS/1607). AG129 mice (male, 5–7 weeks old) and A129 mice (female, 6–8 weeks old) were obtained from an in-house breeding facility at Duke-NUS Vivarium and housed in individually ventilated cages with *ad libitum* food and water. Mouse-adapted DENV-2 S221 strain (kindly gifted by Prof. Subhash Vasudevan, Duke-NUS Medical School) was used for all inoculations.

The DENV antibody-dependent enhancement (ADE) infection model was established as previously described^57,58^. A129 mice were injected with 50 µg 4G2 pan-flavivirus non-neutralizing monoclonal antibody (kindly gifted by Asst. Prof. Satoru Watanabe, Duke-NUS Medical School) one day prior to intravenous (i.v.) inoculation with > 10^7^ plaque-forming units (pfu) virus to induce infection. For the non-ADE severe dengue model, AG129 mice were inoculated i.v. with 10□ pfu virus without 4G2 mAb pre-injection, as reported^59^. Sham-infected controls received sterile PBS.

### Drug treatment

MPO inhibitors AZD5904 and mitiperstat (MTP; AZD4831) (MedChemExpress Singapore) were reconstituted in 100% DMSO per the manufacturer’s protocol. Working solutions were prepared in 10% DMSO, 40% PEG300, 5% Tween-80, and 45% saline were administered *via* oral gavage twice-daily, starting 6 h post-infection, for seven doses. A129 mice received 10 mg/kg per dose, while AG129 mice received 50 mg/kg doses. Vehicle controls were prepared in same solution without inhibitors. Mice were monitored daily and euthanized upon exceeding 20% body weight loss.

### Ex vivo assays

At specified time points post-infections, mice were euthanized and tissues (blood and liver) were harvested. Tissues were either immediately frozen for RNA/protein assays or fixed in 10% neutral buffered formalin for 48-72 h as described^60^ for histopathological analysis. Viral loads were quantified *via* real-time quantitative RT-PCR^61^.

Proteins were extracted from frozen liver homogenates and analyzed for MPO (R&D Systems, DY3667), AST (Abcam, Ab263882), ALT (Abcam, Ab282882), and 4-hydroxynonenal (4-HNE; Abcam, Ab238538) by ELISA following manufacturer’s protocols. Cytokines and chemokines were measured using the Immune Monitoring 48-plex mouse ProcartaPlex panel (Thermo-Fisher, EPX480-20834-901) on a Luminex xMAP platform.

FFPE tissue sections (4 µm) were subjected to immunofluorescence staining using antibodies to probe expression of mouse MPO (Abcam, Ab300650), CD45 (Abcam, Ab10558), neutrophil elastase (NE; Thermo-Scientific, MA5-32548), Ly6G (Abcam, Ab238132), and DNA/RNA damage oxidation products (Abcam, Ab183395). Autofluorescence was quenched using Vector TrueView® (SP-8400-15), and slides were imaged on a Nikon Ni-E inverted fluorescence microscope. Hematoxylin and eosin staining of tissue sections was conducted as previously described^60^.

### Statistical Analysis and Data visualization

Statistical analyses and generation of graphs and heatmaps were performed with Prism v.10.6.1 Software (GraphPad, USA). Categorical variables were compared using χ² tests. Normality of data distribution of continuous data was first determined with Shapiro-Wilk test and QQ plots. Variances among groups of normally distributed data were evaluated with Bartlett’s test. Means from two normally distributed groups were compared with either standard *t*-test or Welch’s *t*-test for groups with equal variances and unequal variances, respectively. Means from > 2 groups of data with normal distribution and equal variances were compared with one-way ANOVA and Tukey’s correction for multiple comparisons. Similarly, means from > 2 groups of data with normal distribution and unequal variances were compared with Welch’s ANOVA and Dunnett’s correction for multiple comparisons. Medians from groups with data that do not follow Normal distribution were compared with either Mann-Whitney for 2 groups, or Kruskal-Wallis test with Dunn’s correction for multiple comparison for > 2 groups.

Kaplan-Meier survival curves were compared using Mantel-Cox test with Holm-Šídák correction for multiple comparisons.

## Acknowledgements

The authors gratefully acknowledge Prof. Subhash Vasudevan and Asst. Prof. Satoru Watanabe from Duke-NUS Medical School, for generously providing the A129 and AG129 mice, mouse-adapted DENV-2 S221 strain, and 4G2 monoclonal antibody used in this study. We also thank Carel Tan Min Jie, Dr. Chaw Su Yin, and Dr. Vanessa Soh from the Duke-NUS Laboratory for Translational and Molecular Imaging for their valuable assistance with *ex vivo* and *in vivo* experiments. We acknowledge Shiau Hui Dong, Diana Tan Bee Har, and Nadiah Binte Abdul Karim for their work and effort in enrolling the patients, and Jaminah D/O Mohamed Ali for human sample processing. We also thank all study participants who volunteered their time and effort to be included in the study.

## Author contributions

A.T., P.Y.C., and T.W.Y. conducted the clinical study in Singapore; P.Y.C. and T.W.Y. performed patient recruitment and sample collection. A.T. conducted the sample processing and plasma analysis; H.K., M.K., D. I., A. W., C. J., and N. M. conducted the clinical study in Sri Lanka and performed the sample collection, processing, and plasma analysis;

K. R., L. L. A., C.A.B.D.O., and R.P.B.D.O conducted the postmortem analysis of fatal dengue cases in Brazil; K.R. and L.L.A. performed the immunohistochemical staining; C.A.B.D.O. and R.P.B.D.O. were responsible for patient recruitment and obtaining informed consent for post□mortem tissue donation; C.B.L.V., S.G., A. G., J.O., J.S.K., and A.-M.C. conducted the animal experiments in Singapore; C.B.L.V. and A-.M.C conceived and designed the animal experiments. C.B.L.V., A.G., S.G., J.O., and J.S.K., performed the animal experiments and *ex vivo* assays.

C.B.L.V., A.T., P. Y. C., and T.W.Y. analysed the data and prepared the figures. T.W.Y. and A.M.-C. supervised; and C.B.L.V. prepared the first draft of the paper, which all authors reviewed.

## Data availability

Raw and analyzed data are available upon request.

