## Supplementary material for "Myeloperoxidase (MPO) exacerbates dengue-associated liver injury and contributes to disease pathogenesis in mouse models": Figure S1

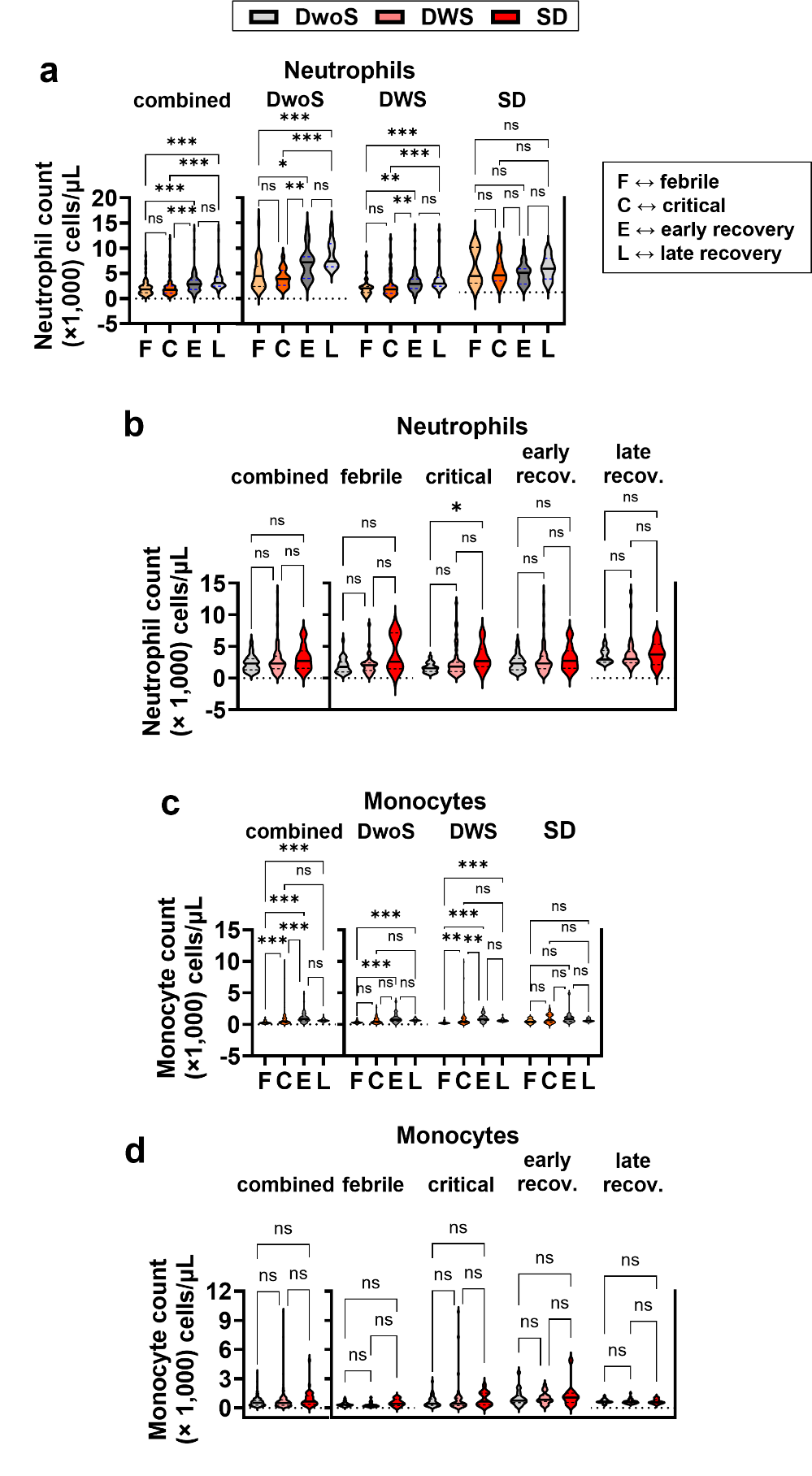

**Figure S1 |** **Temporal changes in blood neutrophil and monocyte counts during dengue illness in Singapore patients.** Dengue cases were classified according to the 2009 WHO criteria: dengue patients without warning signs (*DwoS*), dengue with warning signs (*DWS*), and severe dengue (*SD*). Patients were sampled several times at febrile (F), critical I, early recovery I, and late recovery (L) phases of disease. **a-b**, Neutrophil counts; and **c-d**, Monocyte counts throughout disease and segregated by disease severity, **a,c**; or in DwoS, DWS, and SD patients and segregated by disease phase, **b,d**. Data are shown as median ± interquartile range (IQR) in bold lines and dashed lines, respectively, with medians compared using Kruskal-Wallis test with Dunn’s correction for multiple comparisons. * *p* < 0.05. ** *p* < 0.005. *** *p* < 0.001. *ns*, not significant. *recov*., recovery.
