## Supplementary material for "Myeloperoxidase (MPO) exacerbates dengue-associated liver injury and contributes to disease pathogenesis in mouse models": Figure S2

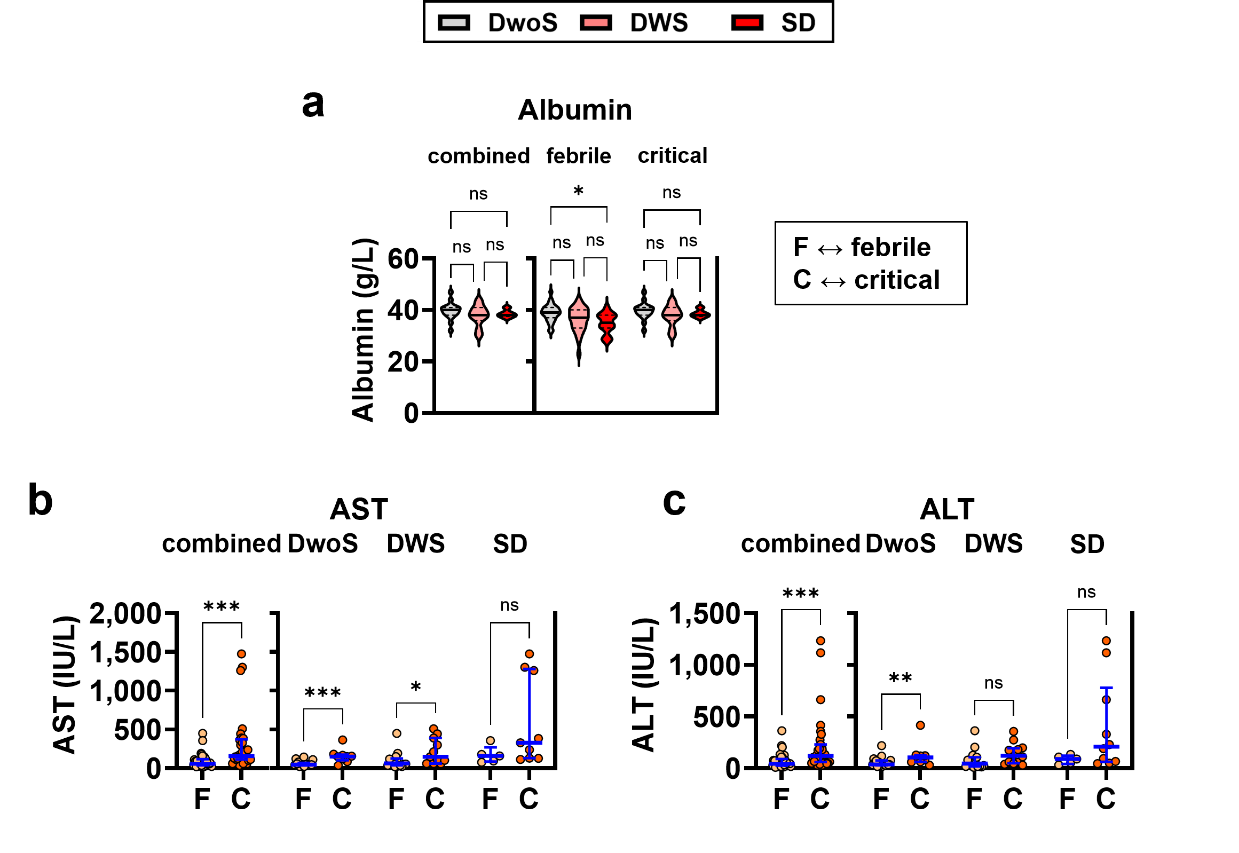

**Figure S2 |** **Plasma albumin and transaminase trajectories across dengue disease phases in Singapore patients.** Dengue cases were classified according to the 2009 WHO criteria: dengue patients without warning signs (DwoS), dengue with warning signs (DWS), and severe dengue (SD). Changes in **a**, albumin; **b**, AST; and **c**, ALT levels at febrile (F) and critical I phases of disease. Data are shown as median ± interquartile range (IQR) values in bold lines and dashed lines, respectively. Medians in ***a*** were compared with Kruskal-Wallis test with Dunn’s correction for multiple comparisons. Medians in ***b-c*** were compared with Mann-Whitney test. * *p* < 0.05. ** *p* < 0.005. *** *p* < 0.001. *ns*, not significant.
