## Supplementary material for "Myeloperoxidase (MPO) exacerbates dengue-associated liver injury and contributes to disease pathogenesis in mouse models": Figure S3

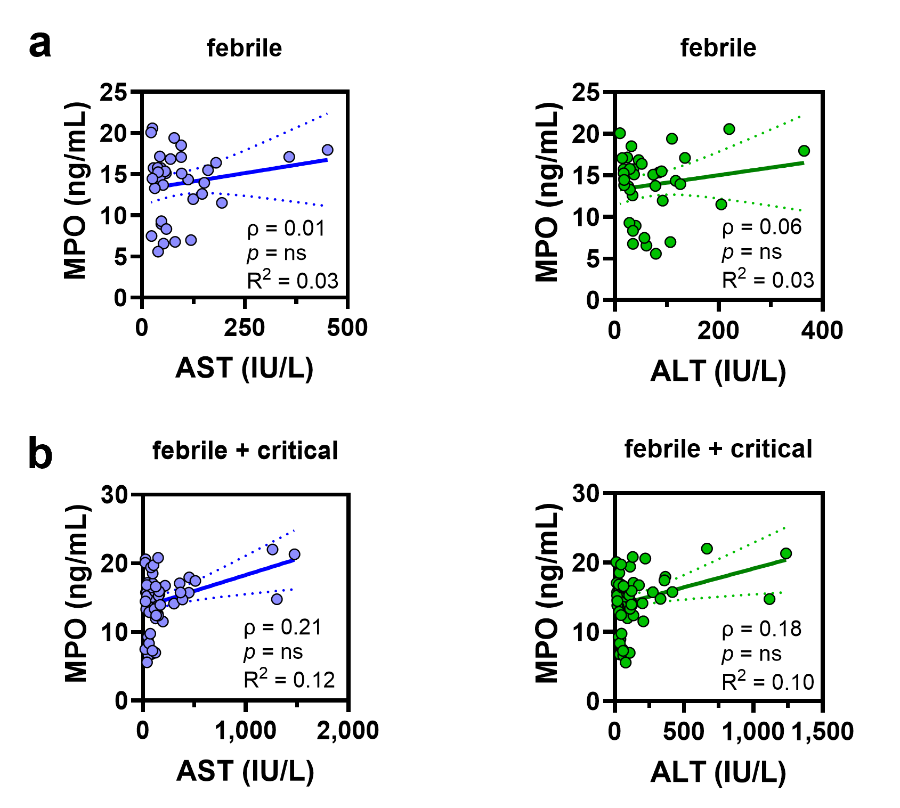

**Figure S3 |** **Lack of correlation between plasma myeloperoxidase (MPO) and liver transaminase in febrile phase of dengue disease in the Singapore cohort.** Monotonic relationship between plasma levels of MPO and either aspartate aminotransferase (AST) or alanine transaminase (ALT) in dengue patients using data from **a**, only febrile phase; or **b**, combining both febrile and critical phases. *Ρ*, Spearman coefficient. *R^2^*, linear coefficient. * *p* < 0.05. ** *p* < 0.005. *** *p* < 0.001. *ns*, not significant.
