## Supplementary material for "Myeloperoxidase (MPO) exacerbates dengue-associated liver injury and contributes to disease pathogenesis in mouse models": Figure S4

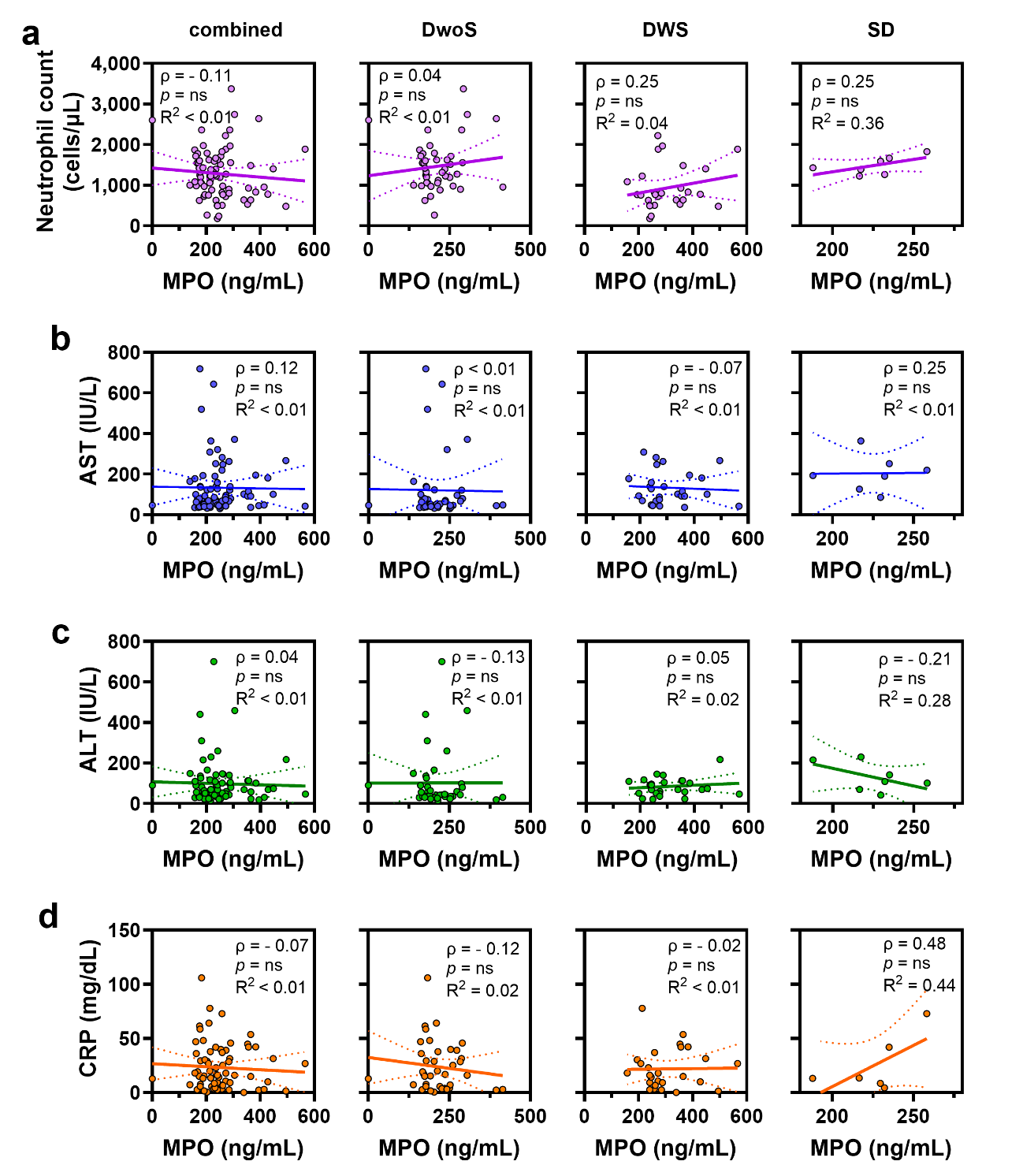

**Figure S4 |** **Lack of correlation between myeloperoxidase (MPO) and markers of neutrophil activation, liver injury, and systemic inflammation in Sri Lanka dengue clinical cohort.** Monotonic relationship between plasma MPO and either **a**, blood neutrophil count; **b**, plasma aspartate aminotransferase (AST); **c**, plasma alanine transaminase (ALT); or **d**, plasma C-reactive protein (CRP) levels. Correlations are shown using data from all patients regardless of disease severity (combined), or from patients segregated into dengue without warning signs (DwoS), dengue with warning signs (DWS), and severe dengue (SD). *ρ*, Spearman coefficient. *R^2^*, linearity coefficient. * *p* < 0.05. ** *p* < 0.005. *** *p* < 0.001. *ns*, not significant.
