## Supplementary material for "Myeloperoxidase (MPO) exacerbates dengue-associated liver injury and contributes to disease pathogenesis in mouse models": Figure S5

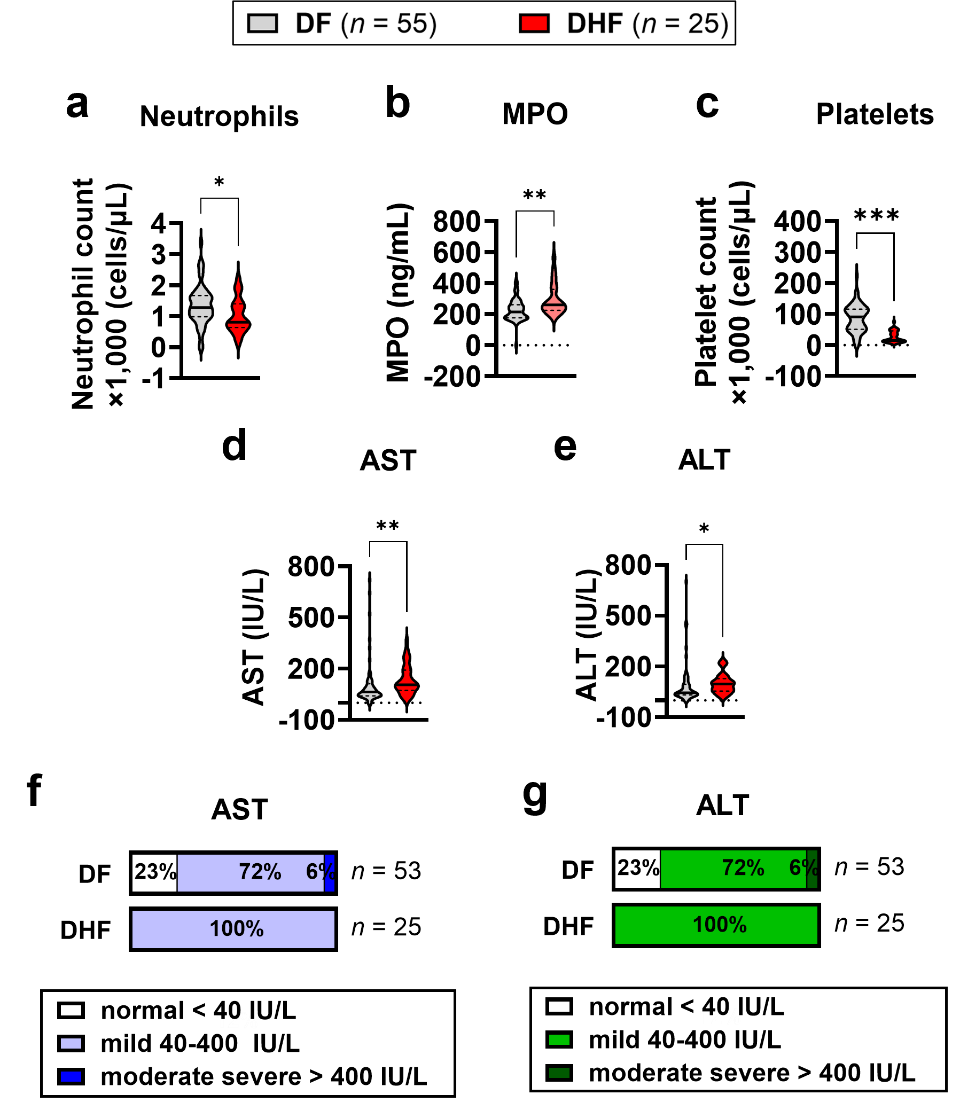

**Figure S5 |** **Circulating myeloperoxidase (MPO) and liver injury markers in the Sri Lanka dengue clinical cohort stratified by 2011 WHO dengue classification.** Dengue patients included in the trial were classified according to 2011 WHO classification as either dengue fever (*DF*), dengue haemorrhagic fever (*DHF*), or dengue shock syndrome (*DSS*). None of the patients exhibited DSS. **A-e** Violin plots depicting median ± interquartile range (IQR) values in bold lines and dashed lines, respectively, of **a**, blood neutrophil counts; and plasma levels of **b**, myeloperoxidase (MPO); **c**, albumin; **d**, aspartate aminotransferase (AST); and **e**, alanine transaminase (ALT). Medians were compared with Mann-Whitney test. **F-g**, Incidence of normal, mild, and moderate severe liver injury based on plasma levels of **f**, AST**;** and **g**, ALT. * *p* < 0.05. ** *p* < 0.005. *** *p* < 0.001. *ns*, not significant.
