## Supplementary material for "Myeloperoxidase (MPO) exacerbates dengue-associated liver injury and contributes to disease pathogenesis in mouse models": Figure S6

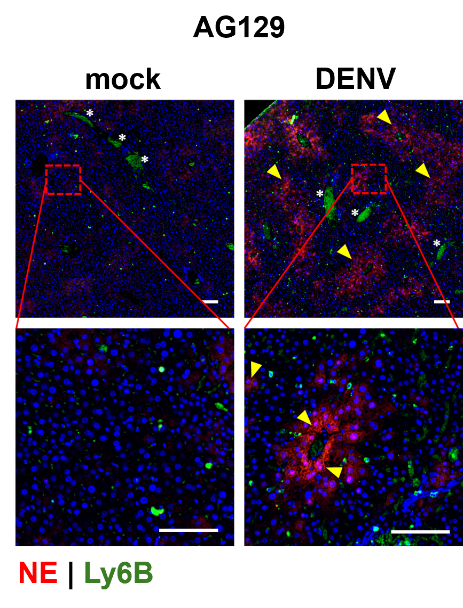

**Figure S6 |** **Activated neutrophils and neutrophil elastase deposition in DENV-infected AG129 mouse livers.** Tissue immunostaining of mouse livers with neutrophil elastase (NE) and myeloid cell marker Ly6B. Areas of NE deposition outside of Ly6B^+^ cells (*yellow arrowheads*), and sites of nonspecific Ly6B staining in liver sinuses (*white asterisks*) are highlighted. Scale bars = 100 µm.
