## Supplementary material for "Myeloperoxidase (MPO) exacerbates dengue-associated liver injury and contributes to disease pathogenesis in mouse models": Figure S7

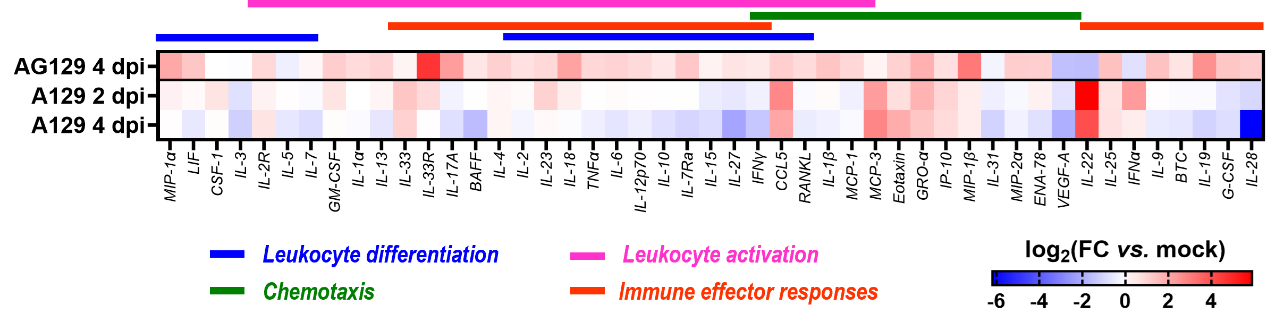

**Figure S7 |** **Induction of broad hepatic inflammatory cytokine and chemokine responses by DENV infection in AG129 and A129 mice.** Heatmap of protein expression in liver homogenates from mice at 2 days and 4 days post-infection (dpi) relative to baseline (mock infection). Protein levels were measured using ProcartaPlex^TM^ 48-plex mouse immune monitoring panel. Chemokines and cytokines involved in leukocyte migration (chemotaxis), differentiation, activation, and immune effector responses are shown. Data are presented as fold-change relative to mock infection (FC *vs.* mock)**.**
