## Supplementary material for "Myeloperoxidase (MPO) exacerbates dengue-associated liver injury and contributes to disease pathogenesis in mouse models": Figure S8

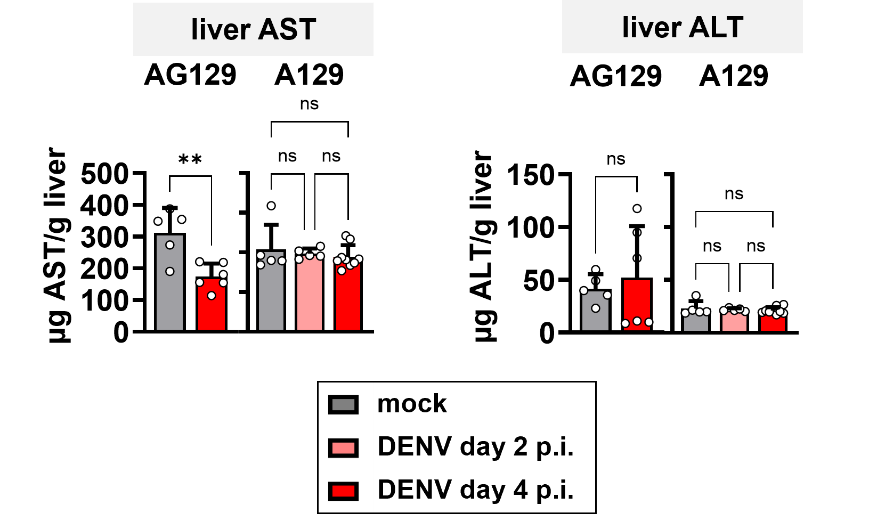

**Figure S8 |** **Hepatic transaminase levels in DENV-infected AG129 and A129 mouse models.** Aspartate aminotransferase (AST) and alanine transaminase (ALT) levels in liver homogenates of AG129 and A129 mice at days 2 and 4 post-infection (p.i.) with dengue virus (DENV). Homogenized livers were diluted > 40,000× to obtain ELISA signals within the acceptable linear range of the standard curve. Data are shown as mean ± SD and compared using either Welch’s *t*-test or Welch’s ANOVA with Dunnett’s correction for multiple comparisons. * *p* < 0.05. ** *p* < 0.005. *** *p* < 0.001. *ns*, not significant.
