## Supplementary material for "Myeloperoxidase (MPO) exacerbates dengue-associated liver injury and contributes to disease pathogenesis in mouse models": Figure S9

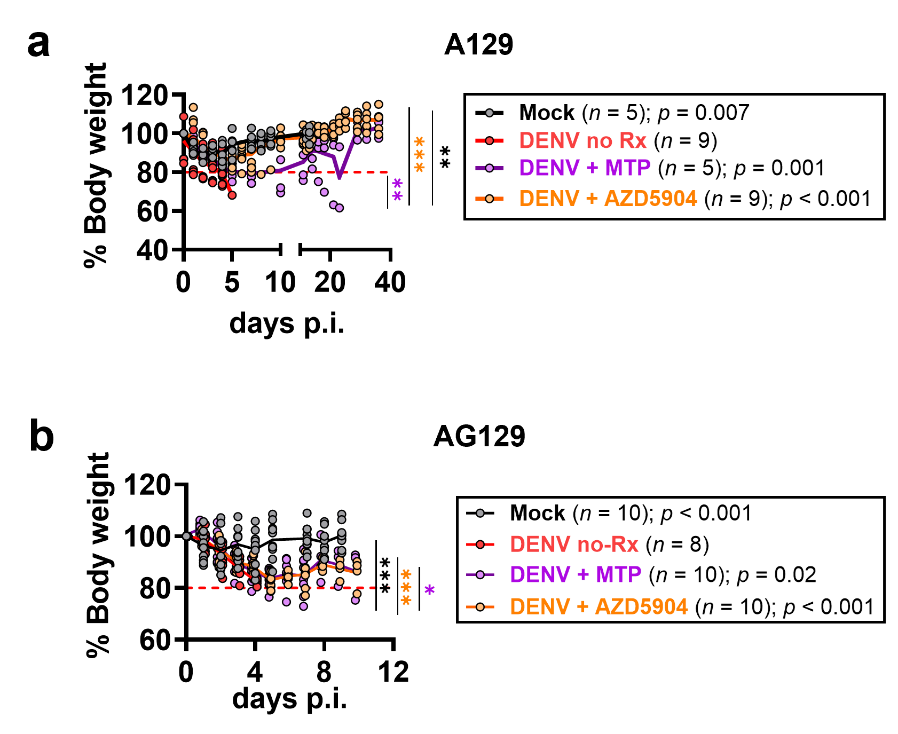

**Figure S9 |** **Rescue of body weight loss in DENV-infected mice following treatment with inhibitors of myeloperoxidase (MPO).** Recorded weights of **a**, A129 mice; and **b**, AG129 mice given twice-daily dose treatments of MPO inhibitors AZD5904 or mitiperstat (MTP). Body weights over time were analysed using area under the curve (AUC) integration analysis. Mean AUCs were compared with Welch’s ANOVA and Dunnett’s correction for multiple comparisons. * *p* < 0.05. ** *p* < 0.005. *** *p* < 0.001. *ns*, not significant.
