## Supplementary material for "Myeloperoxidase (MPO) exacerbates dengue-associated liver injury and contributes to disease pathogenesis in mouse models": Figure S10

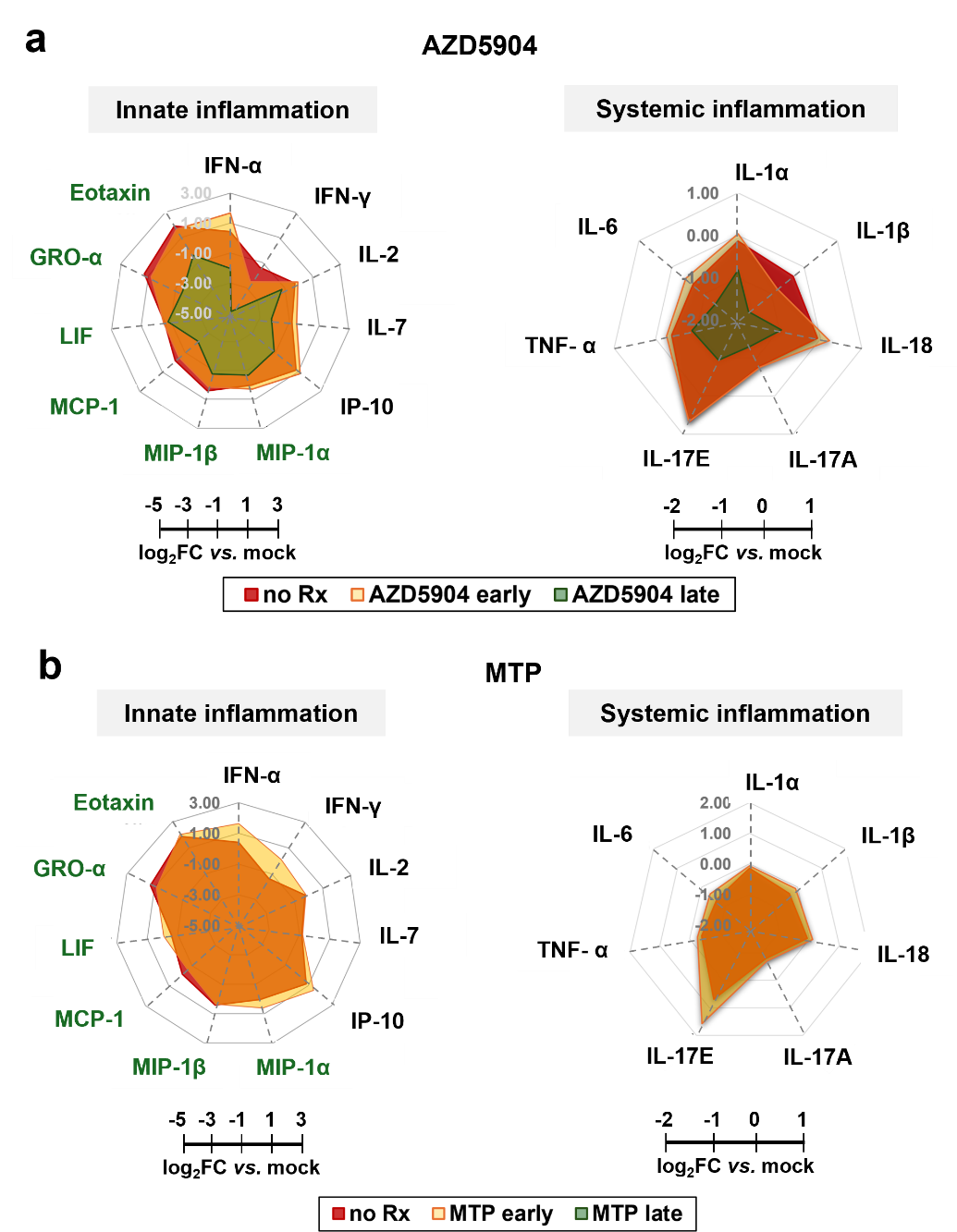

**Figure S10 |** **Myeloperoxidase (MPO) inhibition remodels hepatic innate and systemic inflammatory responses in the A129 DENV antibody-dependent enhancement (ADE) model.** Radial plots of cytokines and chemokines involved in innate inflammation and systemic inflammation from livers of mice treated with either **a**, AZD5904, or **b**, mitiperstat (MTP). Tissues were collected at 4 days (early) or 14 days (late) post-infection with dengue virus (DENV), and tissue lysates were subjected to multiplex ELISA using Immune Monitoring 48-plex mouse ProcartaPlex panel (Thermo-Fisher, EPX480-20834-901) on a Luminex xMAP platform***.*** Cytokine profiles at late treatment are not determined. Proteins involved in neutrophil migration and maturation are shown in green, and data are presented as log_2_ fold-change (log_2_FC) relative to mock infection (FC *vs.* mock).
