## Supplementary material for "Myeloperoxidase (MPO) exacerbates dengue-associated liver injury and contributes to disease pathogenesis in mouse models": Figure S11

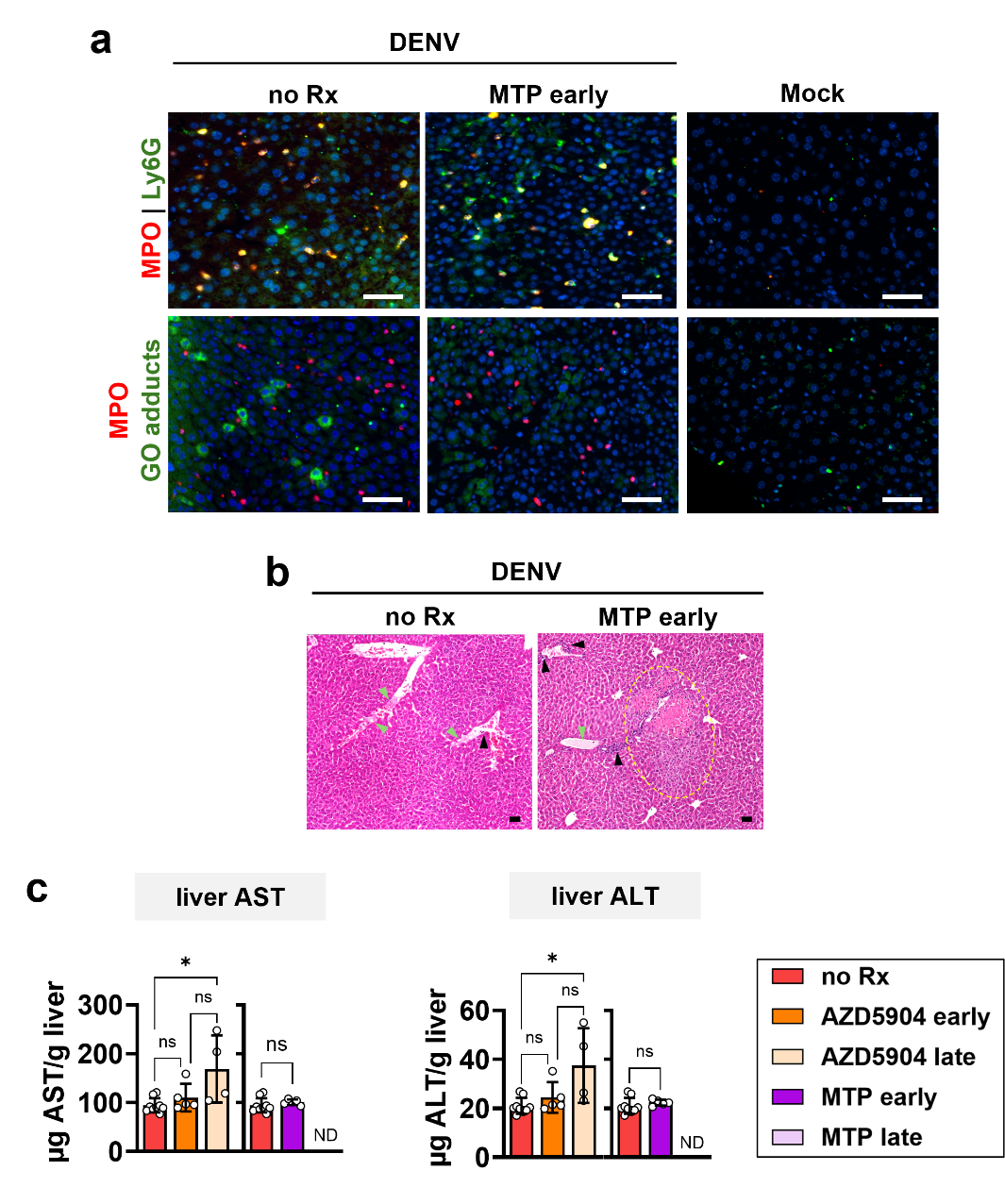

**Figure S11 |** **Liver damage and injury after myeloperoxidase (MPO) inhibition in lethal dengue infection.** **a**, Immunofluorescence staining of liver sections depicting expression of MPO, Ly6G in neutrophils, and guanine-oxidation (GO) products of damaged nucleic acids. Colocalized signals are indicated by white arrows. **b**, Liver sections stained with hematoxylin and eosin. Inflammatory infiltrates are indicated by black arrows, and proteinaceous edema are shown by green arrows. Regions of tissue necrosis are outlined in yellow dashed lines. **c**, Levels of aspartate aminotransferase (AST) and alanine transaminase (ALT) in homogenized livers. Livers were harvested at either 4 days (early) or 14 days (late) post-infection. Data are presented as mean ± s.d., and statistical comparisons between two groups were performed using two-tailed Welch’s *t*-test. Means from three groups were compared with Welch’s ANOVA with Dunnett’s correction for multiple comparison. * *p* < 0.05; ** *p* < 0.005; *** *p* < 0.001; *ns*, not significant. *ND*, not determined. Scale bars = 50 µm.
