## Supplementary material for "Myeloperoxidase (MPO) exacerbates dengue-associated liver injury and contributes to disease pathogenesis in mouse models": Figure S11

**Figure S12**

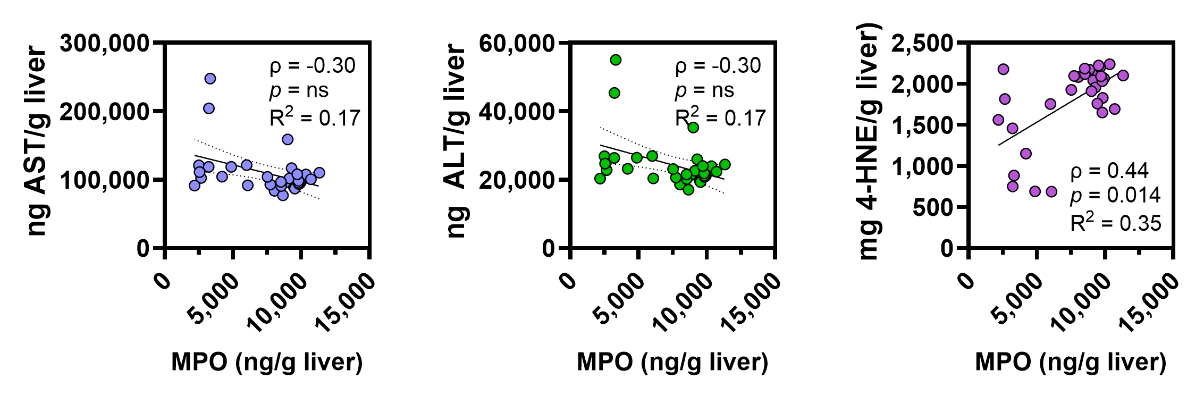

**Figure S11 |** **Monotonic correlation among myeloperoxidase (MPO), aspartate aminotransferase (AST), alanine transferase (ALT), and 4-hydroxynonenal (4-HNE) in DENV-infected livers.** Spearman correlation (*ρ*) between MPO and AST, MPO and ALT, and MPO and 4-HNE in homogenized livers from mice infected with dengue virus (DENV) and subsequently treated with pharmacologic MPO inhibitors AZD5904 or mitiperstat. *R^2^, linear coefficient.*
