## Supplementary material for "Myeloperoxidase (MPO) exacerbates dengue-associated liver injury and contributes to disease pathogenesis in mouse models": Figure S13

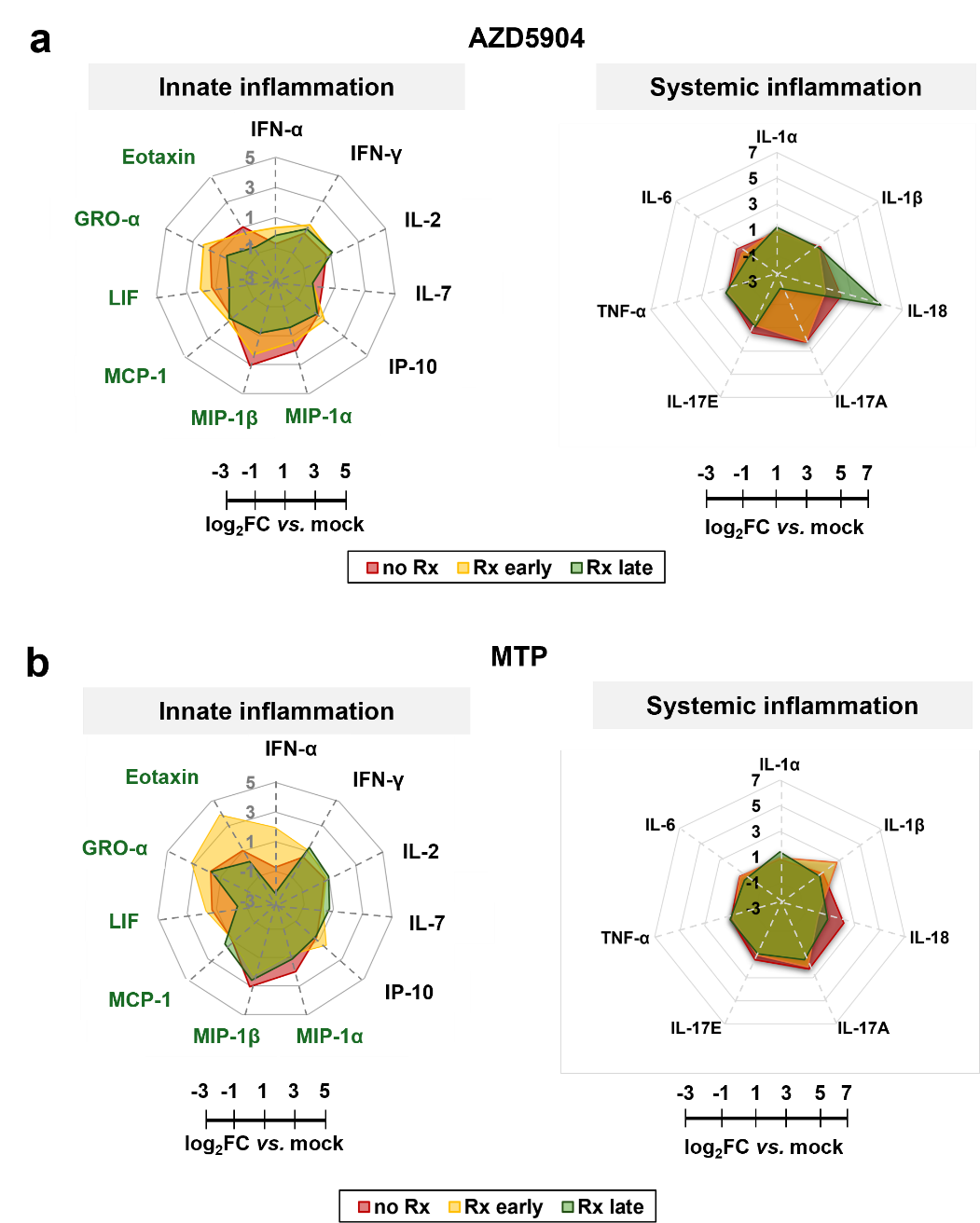

**Figure S13 |** **Innate and systemic inflammatory proteins in AG129 severe dengue model following treatment with inhibitors of myeloperoxidase (MPO).** Radial plots of cytokines and chemokines involved in innate inflammation and systemic inflammation in livers from mice treated with either **a**, AZD5904, or **b**, mitiperstat (MTP). Tissues were collected at 4 days (early) or 8 days (late) post-infection***.*** Proteins involved in neutrophil migration and maturation are shown in green. Data are presented as log_2_ fold-change relative to mock infection (log_2_FC *vs.* mock).
