## Supplementary material for "Myeloperoxidase (MPO) exacerbates dengue-associated liver injury and contributes to disease pathogenesis in mouse models": Figure S14

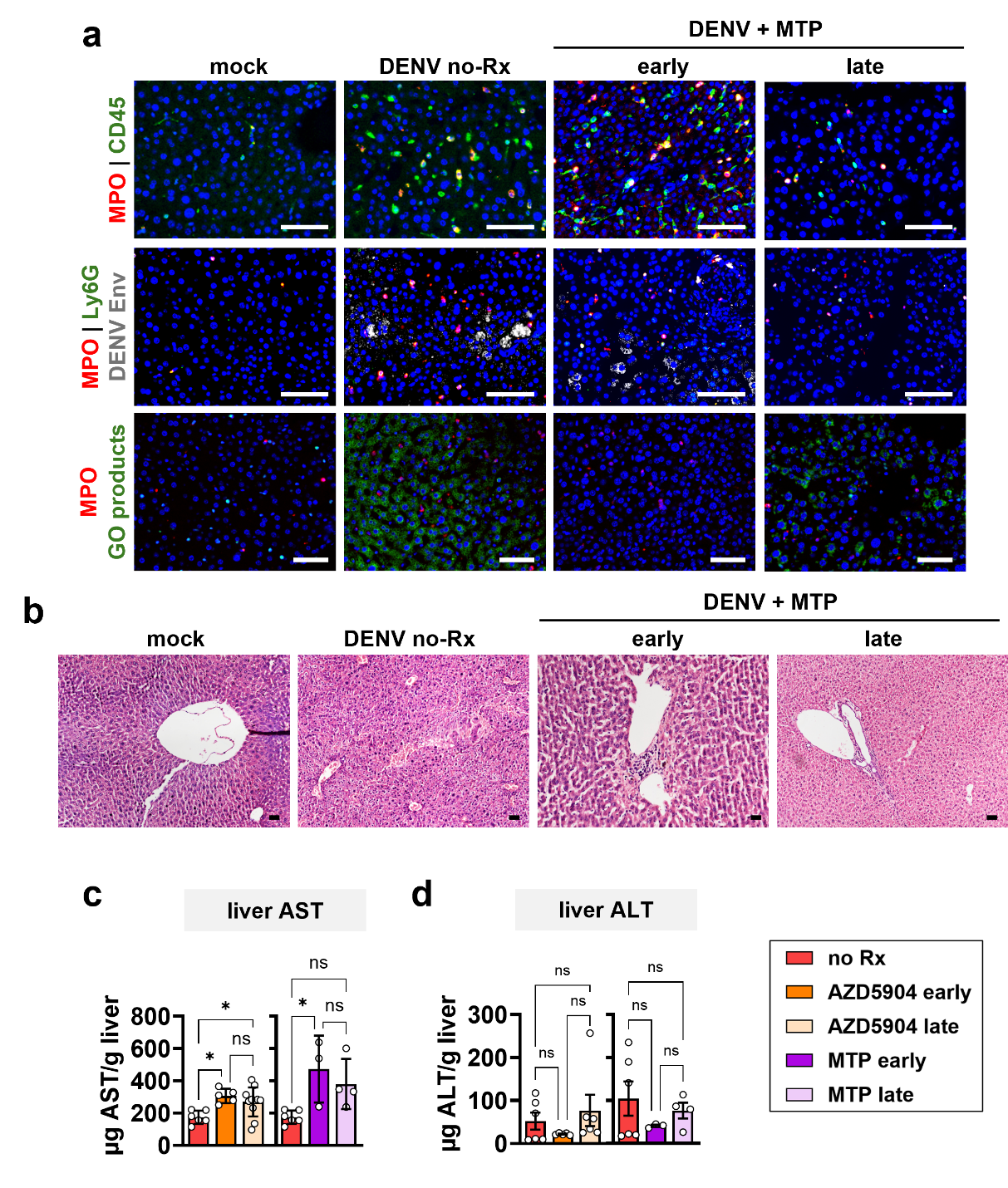

**Figure S14 |** **Liver damage and injury after myeloperoxidase (MPO) inhibition with mitiperstat in a mouse model of DENV infection.** AG129 mice infected with dengue virus (DENV) were treated with mitiperstat (MTP). **a**, Immunofluorescence staining of liver sections depicting expression of MPO, Ly6G in neutrophils, and guanine-oxidation (GO) products of damaged nucleic acids. **b**, Liver sections stained with hematoxylin and eosin. **c-d**, Levels of **c**, aspartate aminotransferase (AST); and **d**, alanine transaminase (ALT) in homogenized livers. Livers were harvested at either 4 days (early) or 8 days (late) post-infection. Data are presented as mean ± s.d. and statistical comparisons were performed using Welch’s ANOVA with Dunnett’s correction for multiple comparisons. * *p* < 0.05; ** *p* < 0.005; *** *p* < 0.001; *ns*, not significant. Scale bars = 50 µm.
