## Supplemental Methods for "Myeloperoxidase (MPO) exacerbates dengue-associated liver injury and contributes to disease pathogenesis in mouse models"

**Singapore dengue study**

Dengue subjects were enrolled in a prospective longitudinal study between September 2017 and December 2019 and were recruited in the febrile or critical phase and followed up at 21-28 d after discharge. Dengue infection was confirmed by a positive NS1 antigen test (SD Bioline Dengue Duo, Korea) and pregnant or breastfeeding patients were excluded. Disease severity was defined according to the WHO 2009 criteria (dengue fever without warning signs (DwoS), dengue fever with warning signs (DWS), and severe dengue (SD)). Liver disease severity was classified according to criteria proposed by a consensus of dengue researchers developing standardized clinical endpoints for interventional trials. Briefly, moderate severe liver disease was defined as i) acute illness with signs and symptoms of acute hepatitis, ii) alanine transaminase (ALT) >10 times upper limit of normal (40 U/L), iii) no criteria for acute liver failure; and severe liver disease/acute liver failure was i) as above with ii) acute mental status change consistent with encephalopathy, and iii) new onset coagulopathy with INR ≥1.5. ALT <40 U/L was defined as normal and mild liver dysfunction was ALT levels >40 U/L and <400 U/L.

In addition to disease severity, demographic, clinical and laboratory information were collected. The critical phase was defined according to the day with the lowest platelet count concurrent with the highest hematocrit and defervescence. Controls were adults with no febrile episodes two weeks prior to recruitment and no previous dengue within the past six months.

Leukocyte counts, neutrophil and monocyte percentages, liver enzymes and albumin levels were obtained from hospital results with analysis being done at the Tan Tock Seng Hospital Clinical Pathology Laboratory. Absolute neutrophil and monocyte counts were calculated as percentage of neutrophil/monocytes x leukocyte counts. Blood for research assays was collected in either a Li-Heparin or sodium citrate tube and spun down within 45 min of collection in a centrifuge and the plasma frozen down at -80°C.

**Sri Lanka dengue study**

A total of 577 adult patients presenting with a suspected acute Dengue infection were recruited from the National Institute of Infectious Diseases, Sri Lanka, between December 2022 and December 2024, following informed written consent. They were considered to have acute Dengue infection if they had a positive NS1 antigen test (SD Biosensor, South Korea) or a positive DENV-specific real-time polymerase chain reaction test. All those with known chronic kidney disease or chronic liver disease were excluded from the study. Data from randomly selected seventy-eight (78) patients are reported in this study.

Patients were recruited during the febrile phase (≤4 d since onset of illness), with the first day of onset of fever considered the first day of illness. Liver transaminases and C-reactive protein (CRP) were measured in all patients on the day of admission (on the day of recruitment into the study). Clinical symptoms and laboratory parameters were recorded multiple times per day while in the hospital. Fluid leakage was assessed daily using bed-side ultrasound scans to detect pleural effusions and ascites. Disease severity was originally classified according to the 2011 World Health Organization (WHO) Dengue classification criteria. Accordingly, patients with a rise in hematocrit of ≥20% from baseline or those with ultrasound evidence of plasma leakage were classified as having DHF. Patients who developed shock, defined by a pulse pressure narrowing to 20 mmHg, were categorized as having Dengue shock syndrome (DSS). For the purpose of this study, patients were reclassified according to the 2009 WHO Guidelines to facilitate direct comparison with the Singapore cohort.

**Clinical history of liver samples from Brazil dengue post-mortem study**

**Case 1 –** A 38-year-old female was brought to the emergency department of Antônio Pedro Hospital after being found unwell in a public area. Upon admission, she presented with bilateral miosis that rapidly progressed to cardiorespiratory arrest and mydriasis. Autopsy findings revealed subarachnoid and brainstem hemorrhages, cerebral edema, pulmonary emphysema with anthracosis, and marked vascular congestion in both spleen and liver.

**Case 2 –** A 21-year-old obese female experienced fever, myalgia, and headache for eight days, followed by metrorrhagia, nausea, abdominal pain, vomiting, and diarrhea. Laboratory tests showed severe leukopenia and thrombocytopenia (platelet count: 10,000/mm³). She was admitted to the ICU of Clementino Fraga Filho University Hospital with respiratory failure. Biochemical analysis indicated hyperglycemia (158 mg/dL), elevated AST (149 IU/L) and ALT (66 IU/L). Abdominal ultrasonography revealed peripancreatic fluid accumulation. The patient’s condition deteriorated to refractory shock and multiple organ failure, resulting in death. Autopsy revealed alveolar edema in lungs, hepatic steatosis and periportal congestion in the liver, and splenic congestion^1^.

**Case 3–** A 41-year-old woman was admitted to Miguel Couto Hospital reporting weakness, syncope, sweating, epigastric pain, fever of two days’ duration, abdominal pain, and yellowish discharge. Laboratory findings included leukocytosis and elevated hematocrit (48%). Abdominal edema was also noted. Death was attributed to visceral congestion and acute pulmonary edema. RT-PCR for DENV-3, as described by Lanciotti *et. al*.^2^, was positive in fresh liver tissue samples. Autopsy revealed pulmonary edema, cardiomyopathy, ischemia, and luteal cysts^1^.

**Case 4 –** A 61-year-old female hospitalized at Miguel Couto Hospital presented with fever, myalgia, vomiting, and diarrhea. Autopsy findings demonstrated hypertrophic cardiomyopathy, peripheral cyanosis, and pericardial effusion. The immediate cause of death was acute pulmonary edema secondary to sudden cardiorespiratory arrest. Autopsy revealed pulmonary edema, cardiomyopathy, hepatic congestion, pyelonephritis, and renal detention^1^.

**Immunostaining of postmortem dengue liver sections.** Detection of specific cell populations and inflammatory markers was performed by immunohistochemistry using the Vector IHQ commercial kit (Vector Laboratories, USA). Tissue sections were incubated with 3% hydrogen peroxide (H_2_O_2_) for 15 min to block endogenous peroxidase activity, followed by three washes in phosphate-buffered saline (PBS, pH 7.4) for 5 min each. Antigen retrieval was carried out in sodium citrate buffer (pH 6.0) or Tris-EDTA buffer (pH 9.0) for 20 min at 60 °C. After cooling to room temperature, sections were incubated with 3% bovine serum albumin (BSA) in PBS for 20 min to reduce nonspecific binding. Subsequently, slides were incubated overnight at 4 °C with the following primary antibodies: rat anti-human MPO monoclonal antibody (Abcam, AB300650; 1:3000), and rabbit anti-human CD177 and monoclonal antibody (Abcam, AB255296; 1:500). After primary antibody incubation, sections were rinsed three times in PBS and incubated with the appropriate secondary antibody (Vector Laboratories, USA) for 30 min at room temperature. The immunoreactive sites were visualized using diaminobenzidine (DAB; Vector Laboratories, USA) as the chromogenic substrate. Slides were counterstained with Harris hematoxylin (Sigma, MO, USA), dehydrated, cleared, and mounted using Entellan® mounting medium. Immunoperoxidase staining was examined under a light microscope equipped with a CCD camera (Olympus BX53 with DP72 camera, Japan). Images were acquired and analyzed using the Image-Pro Plus 7.0 software (Media Cybernetics, USA).
