## Supplemental Tables for "Myeloperoxidase (MPO) exacerbates dengue-associated liver injury and contributes to disease pathogenesis in mouse models"

**Table S1 |** Blood parameters and relevant plasma protein levels in Singapore dengue patients sampled multiple times during disease and recovery. Data are shown as median (interquartile range, IQR) values compared among dengue without warning signs (DwoS), dengue with warning signs (DWS), and severe dengue (SD) patients.

| Variables | Controls | DwoS | DWS | SD | *p*^a^ |
| --- | --- | --- | --- | --- | --- |
| Absolute Neutrophil Counts (cells/µL) | | | | | |
| Combined^b^ | NA***^c^*** | 2,319  (1,369-3,075) | 2,322  (1,496-3,477) | 2,739  (1,571-4,284) | 0.07 |
| Febrile Phase | NA | 1,795  (1,001-2,457) | 2,036  (1,224-2,412) | 2,595  (1,571-6,864) | 0.26 |
| Critical Phase |  | 1,560  (1,067-2,106) | 1,825  (1,073-2,496) | 2,700  (2,112-4,284) | **0.02** |
| Early Recovery Phase |  | 2,883  (1,645-3,265) | 2,060  (1,595-3,468) | 2,456  (1,315-3,502) | 0.07 |
| Late Recovery Phase |  | 2,945  (2,561-4,265) | 3,002  (2,466-4,179) | 3,757  (2,717-4,845) | 0.84 |
| Absolute Monocyte Count (cells/µL ) | | | | | |
| Combined^b^ | NA | 534  (307-781) | 512  (279-828) | 662  (391-1,214) | 0.06 |
| Febrile Phase | NA | 308  (222-422) | 219  (176-317) | 419  (391-684) | 0.07 |
| Critical Phase |  | 444  (261-911) | 440  (265-1,001) | 654  (352-1,520) | 0.44 |
| Early Recovery Phase |  | 761  (547-1,154) | 1,269  (715-2,466) | 1,073  (634-1,335) | 0.70 |
| Late Recovery Phase |  | 636  (575-776) | 597  (503-710) | 577  (518-748) | 0.44 |
| Myeloperoxidase, MPO (ng/mL) | | | | | |
| Combined^b^ | 8.83  (4.45-14.81) | 13.73  (11.42-15.76) | 14.11  (12.38-16.80) | 14.75  (13.14-20.89) | **< 0.001** |
| Febrile Phase |  | 13.73  (8.83-15.75) | 13.81  (11.51-15.73) | 21.6  (18.50-22.52) | **0.006** |
| Critical Phase |  | 14.15  (11.56-15.75) | 14.49  (13.2-16.87) | 15.91  (13.78-21.10) | 0.10 |
| Early Recovery Phase |  | 12.65  (11.54-13.19) | 14.48  (12.50-16.57) | 12.79  (12.35-13.23) | 0.20 |
| Late Recovery Phase |  | 14.91  (12.17-16.61) | 13.34  (7.94-15.99) | 13.83  (9.46-14.17) | 0.24 |
| Albumin (g/L) | | | | | |
| Combined^b^ |  | 39  (37-41) | 37  (34-40) | 35  (33-38) | 0.31 |
| Febrile Phase | NA | 40  (38-41) | 38  (37-41) | 38  (38-39) | **0.01** |
| Critical Phase |  | 37  (36-39) | 35  (33-38) | 34  (30-35) | 0.31 |
| Alanine transaminase, ALT (IU/L) | | | | | |
| Combined^b^ |  | 61  (35-98) | 74  (30-154) | 110  (59-280) | **0.04** |
| Febrile Phase | NA | 37  (30-70) | 46  (18-96) | 90  (52-110) | 0.37 |
| Critical Phase |  | 104  (63-122) | 123  (56-186) | 210  (73-581) | 0.30 |
| Aspartate aminotransferase, AST (IU/L) | | | | | |
| Combined^b^ |  | 59  (43-141) | 96  (56-194) | 210  (130-378) | **0.001** |
| Febrile Phase | NA | 47  (38-59) | 63  (39-113) | 160  (95-180) | **0.03** |
| Critical Phase |  | 152  (123-169) | 147  (82-347) | 330  (131-1,261) | 0.15 |

*^a^* Kruskal-Wallis test for comparison of controls, dengue without warning signs (DwoS), dengue with warning signs (DWS), and severe dengue (SD) groups.

*^b^* Combined data refers to all dengue patients of varying severity regardless of disease phase (febrile, critical or recovered).

***^c^*** NA, not applicable.

**Table S2** | Blood parameters and relevant plasma protein levels in Singapore dengue patients sampled multiple times during disease and recovery. Data are shown as median (interquartile range, IQR) values compared during febrile, critical, and recovery phases of disease.

| Variables | Controls | | Febrile | Critical | Early  recovery | Late  recovery | *p*^a^ |
| --- | --- | --- | --- | --- | --- | --- | --- |
|  | | **Absolute Neutrophil Counts (cells / µL)** | | | | | |
| Combined^b^ | NA***^c^*** | | 1,833  (1,172-2,499) | 1,756  (1,140-2,483) | 2,868  (1,858-3,701) | 3,087  (2,502-4,252) | **< 0.001** |
| DwoS | NA | | 1,795  (1,001-2,457) | 1,560  (1,067-2,106) | 2,883  (1,645-3,265) | 2,945  (2,561-4,265) | **< 0.001** |
| DWS |  |  | 2,036  (1,224-2,412) | 1,825  (1,073-2,496) | 2,942  (2,036-3,918) | 3,002  (2,466-4,179) | **< 0.001** |
| SD |  |  | 2,595  (1,571-6,864) | 2,700  (2,112-4,284) | 2,456  (1,315-3,502) | 3,757  (2,717-4,845) | 0.72 |
|  | | **Absolute Monocyte Counts (cells / µL)** | | | | | |
| Combined^b^ | NA | | 283  (196-418) | 467  (265-952) | 862  (585-1,242) | 615  (530-747) | **< 0.001** |
| DwoS | NA | | 308  (222-422) | 444  (261-911) | 761  (547-1,154) | 636  (575-776) | **< 0.001** |
| DWS |  |  | 219  (176-317) | 440  (265-1,001) | 840  (668-1,232) | 597  (503-710) | **< 0.001** |
| SD |  |  | 419  (391-684) | 654  (352-1,520) | 1,073  (634-1,335) | 577  (518-748) | 0.51 |
|  | | **Myeloperoxidase, MPO (ng/mL)** | | | | | |
| Combined^b^ | 8.83  (4.58-14.81) | | 13.93  (10.46-17.00) | 14.49  (12.83-16.53) | 13.42  (12.33-15.18) | 13.81  (10.34-15.56) | **0.002** |
| DwoS | NA | | 13.73  (8.83-15.75) | 14.14  (11.56-15.79) | 12.64  (11.54-13.18) | 14.91  (12.17-16.61) | 0.56 |
| DWS |  |  | 13.81  (11.51-15.73) | 14.49  (13.22-16.87) | 14.47  (12.50-16.57) | 13.34  (7.94-15.99) | 0.17 |
| SD |  |  | 21.60  (18.50-22.52) | 15.91  (13.78-21.10) | 12.79  (12.35-13.23) | 13.83  (9.46-14.17) | **0.01** |
|  | | **Alanine transaminase, ALT (IU/L)** | | | | | |
| Combined^b^ | NA | | 43  (28-89) | 120  (62-215) | NA | NA | **< 0.001** |
| DwoS | NA | | 37  (30-70) | 104  (63-122) | NA | NA | **0.009** |
| DWS |  |  | 46  (18-96) | 123  (56-186) | NA | NA | 0.05 |
| SD |  |  | 90  (52-110) | 210  (73-581) | NA | NA | 0.16 |
|  | | **Aspartate aminotransferase, AST (IU/L)** | | | | | |
| Combined^b^ | NA | | 57  (39-114) | 156  (115-366) | NA | NA | **< 0.001** |
| DwoS | NA | | 47  (38-59) | 152  (123-169) | NA | NA | **< 0.001** |
| DWS |  |  | 63  (39-113) | 147  (82-347) | NA | NA | **0.02** |
| SD |  |  | 160  (95-180) | 330  (131-1,261) | NA | NA | 0.15 |
|  | | **Albumin (g/L)** | | | | | |
| Combined^b^ | NA | | 39  (37-41) | 35  (33-38) | NA | NA | **< 0.001** |
| DwoS | NA | | 40  (38-41) | 37  (36-39) | NA | NA | **0.04** |
| DWS |  |  | 38  (37-41) | 35  (33-38) | NA | NA | 0.10 |
| SD |  |  | 38  (38-39) | 34  (30-35) | NA | NA | **0.004** |

*^a^* Mann-Whitney or Kruskal-Wallis test for comparison of febrile, critical, and recovered dengue patients.

*^b^* Combined data refers to all febrile, critical or recovered patients regardless of disease severity (DwoS, DWS or SD).

***^c^*** NA, not applicable.

**Table S3 |** Blood cell counts and plasma levels of relevant proteins in dengue patients from Sri Lanka classified using WHO 2009 dengue guidelines. Data are shown as median (interquartile range, IQR).

| Variables |  | DwoS  (n = 44) | DWS  (n = 27) | SD  (n = 7) | *p*^a^ |
| --- | --- | --- | --- | --- | --- |
| Lowest neutrophil count (cells/µL) |  | 1,200  (1,060-1,710) | 770  (580-1,160) | 1,430  (1,330-1,630) | **< 0.001** |
| Lowest leukocyte count (cells/µL) |  | 2,850  (2,490-3,610) | 2,860  (2,160-3,320) | 2,880  (2,680-3,050) | 0.68 |
| Lowest platelet count (cells/µL) |  | 101,000  (66,300-118,300) | 30,000  (14,500-44,500) | 45,000  (13,000-62,000) | **< 0.001** |
| Myeloperoxidase, MPO (ng/mL) |  | 197.7  (176.2-251.2) | 271.6  (241.4-360.6) | 229.5  (216.9-233.5) | **< 0.001** |
| Aspartate aminotransferase, AST (IU/L) |  | 61  (43-100) | 102  (72-180) | 194  (158-236) | **0.002** |
| Alanine transaminase, ALT (IU/L) |  | 42  (34-90) | 74  (51-111) | 109  (85-179) | **0.03** |
| C-reactive protein, CRP (mg/dL) |  | 16.7  (5.66-36.9) | 17.3  (9.2-30.8) | 13.3  (9.6-34.8) | 0.99 |

*^a^* Kruskal-Wallis test for comparison of DwoS, DWS, and SD patients.

**Table S4** | Baseline characteristics of the Sri Lanka dengue participants by 2011 Dengue guidelines. Patients were classified as having dengue fever (DF), dengue haemorrhagic fever (DHF), or dengue shock syndrome (DSS). None of the patients exhibited DSS. Data are shown as median (interquartile range, IQR).

| Variables | DF  (n = 53) | DHF  (n = 25) | *p*^a^ |
| --- | --- | --- | --- |
| Male (%) | 35 (66.0) | 19 (76.0) | 0.455 |
| Median age (IQR) [range], years | 35  (28-46) | 32  (21-40) | 0.412 |
| Median Body Mass Index (IQR), kg/m^2^ | 24.4  (20.8-26.4) | 24.1  (21.6-28.2) | 0.45 |
| Diabetes Mellitus, n (%) | 5 (9.1) | 1 (4.0) | 0.88 |
| Hypertension, n (%) | 4 (7.3) | 1 (4.0) | 0.89 |

*^a^* Mann-Whitney or chi-squared test for comparison of dengue fever (DF) and dengue haemorrhagic fever (DHF) groups.

**Table S5 |** Blood cell counts and plasma levels of relevant proteins in dengue patients from Sri Lanka classified using WHO 2011 dengue guidelines. Patients were classified as having dengue fever (DF), dengue hemorrhagic fever (DHF), or dengue shock syndrome (DSS). None of the patients exhibited DSS. Data are shown as median (interquartile range, IQR).

| Variables |  | DF  (n = 53) | DHF  (n = 25) | *p*^a^ |
| --- | --- | --- | --- | --- |
| Lowest neutrophil count ×1,000 (cells/µL) |  | 1.27  (0.98-1.67) | 0.80  (0.64-1.39) | **0.01** |
| Lowest leukocyte count ×1,000 (cells/µL) |  | 2.88  (2.30-3.40) | 2.88  (2.38-3.33) | 0.83 |
| Lowest platelet count ×1,000 (cells/µL) |  | 96.0  (54.0-116.0) | 21.0  (13.0-39.0) | **< 0.001** |
| Myeloperoxidase, MPO (ng/mL) |  | 220.5  (177.6-260.9) | 259.3  (232.0-356.4) | **0.001** |
| Aspartate aminotransferase, AST (IU/L) |  | 63  (41-101) | 139  (92-220) | **0.004** |
| Alanine transaminase, ALT (IU/L) |  | 44  (31-91) | 96  (58-117) | **0.01** |
| C-reactive protein, CRP (mg/dL) |  | 15.70  (3.50-36.42) | 14.89  (9.37-30.30) | 0.46 |

*^a^* Mann-Whitney test for comparison of DF *vs.* DHF patients.

**Table S6 |** Liver immunohistochemistry in Brazil fatal dengue samples. Staining data for MPO and CD177 are shown as median (interquartile range, IQR), and values are compared among four dengue positive cases and non-infectious control.

|  | Control | Case 1 | | Case 2 | | Case 3 | | Case 4 |
| --- | --- | --- | --- | --- | --- | --- | --- | --- |
|  | **MPO** | | | | | | | |
| Median pixels/area  (IQR*^a^*)  *p^b^* | 103  (74-155)  - | | 3,021  (2,650-3,288)  <0.001 | | 2,536  (1,660-3,692)  0.001 | 2,707  (2,457-3,088)  <0.001 | 122  (88-237)  0.8 | |
| Fold-change |  | | 29.4 | | 24.7 | 26.4 | 1.2 | |
|  | **CD177** | | | | | | | |
| Median pixels/area  (IQR*^a^*)  *p^b^* | 376  (358-404)  - | 875  (721-932)  <0.001 | | 517  (421-816)  0.13 | | 598  (536-780)  0.14 | | 956  (739-1,267)  <0.001 |
| Fold-change |  | 2.3 | | 1.4 | | 1.6 | | 2.5 |

*^a^* IQR; interquartile range

*^b^* Kruskal-Wallis test with Dunn’s correction for multiple-comparison among groups
